# Developmental cell fate choice employs two distinct cis regulatory strategies

**DOI:** 10.1101/2022.06.06.494792

**Authors:** M. Joaquina Delás, Christos M Kalaitzis, Tamara Fawzi, Madeleine Demuth, Isabel Zhang, Hannah T Stuart, Elena Costantini, Kenzo Ivanovitch, Elly M Tanaka, James Briscoe

## Abstract

In many developing tissues the patterns of gene expression that assign cell fate are organised by secreted signals functioning in a graded manner over multiple cell diameters. Cis Regulatory Elements (CREs) interpret these graded inputs to control gene expression. How this is accomplished remains poorly understood. In the neural tube, a gradient of the morphogen Sonic hedgehog allocates neural progenitor identity. Here, we uncover two distinct ways in which CREs translate graded Shh signaling into differential gene expression. In the majority of ventral neural progenitors a common set of CREs are used to control gene activity. These CREs integrate cell type specific inputs to control gene expression. By contrast, the most ventral progenitors use a unique set of CREs. These are established by the pioneer factor FOXA2, paralleling the role of FOXA2 in endoderm. Moreover, FOXA2 binds a subset of the same sites in neural and endoderm cells. Together the data identify distinct cis regulatory strategies for the interpretation of morphogen signaling and raise the possibility of an evolutionarily conserved role for FOXA2-mediated regulatory strategy across tissues.

## INTRODUCTION

During development signalling cues direct cell fate decisions. Precise spatiotemporal gene expression is essential for this process. Cis Regulatory Elements (CREs) integrate inputs from tissue and cell type specific transcription factors (TFs), as well as signalling effectors, to direct gene expression (Spitz and Furlong, 2012). This suggests a straightforward mechanism in which distinct signals produce different cell types by activating different transcriptional effectors. However, in many tissues a single signal directs multiple, alternative cell fates (Rogers and Schier, 2011). Two strategies can be envisioned for how CREs interpret a single signal to define multiple cell fates. Different CREs could function in different cell types. In this strategy – Differential Accessiblity – the availability of an element is the principal determinant of cell identity. In an alternative strategy – Differential Binding – the same CREs could be used in all cell types in the tissue and the configuration of proteins bound to the elements determines cell fate. An implication of these two strategies is that choosing between two fates will require chromatin remodelling in the case of the differential availability of CREs but can happen without remodelling in the case of different protein configurations at the CREs. A well characterised example of a signal controlling multiple cell types is the morphogen Sonic Hedgehog (Shh). Shh controls the pattern of neuronal subtype generation in ventral regions of the developing neural tube (Jessell, 2000). Shh, initially secreted from the notochord and later from the floor plate (Echelard et al., 1993; Roelink et al., 1994), forms a ventral to dorsal gradient in the neural tube and establishes a set of molecularly distinct domains that occupy characteristic positions along the dorsal-ventral axis. Progenitors in each domain differentiate into distinct classes of post-mitotic neurons. This organisation is necessary for the subsequent assembly of the locomotor and sensory circuits (Balaskas et al., 2019; Bikoff et al., 2016). The progenitor domains are identifiable by characteristic gene expression programmes (Fig 1A). NKX2.2 is expressed in p3 progenitors, the most ventral domain (Briscoe et al., 1999); OLIG2 is expressed in adjacent motor neuron progenitors (pMN) (Novitch et al., 2001); PAX6 is expressed in more dorsal progenitor domains, including p0, p1 and p2 domains while lower expression is detected in pMN (Ericson et al., 1997); NKX6.1 is expressed in, p3, pM and p2 domains (Briscoe et al., 2000).

**FIG. 1:**
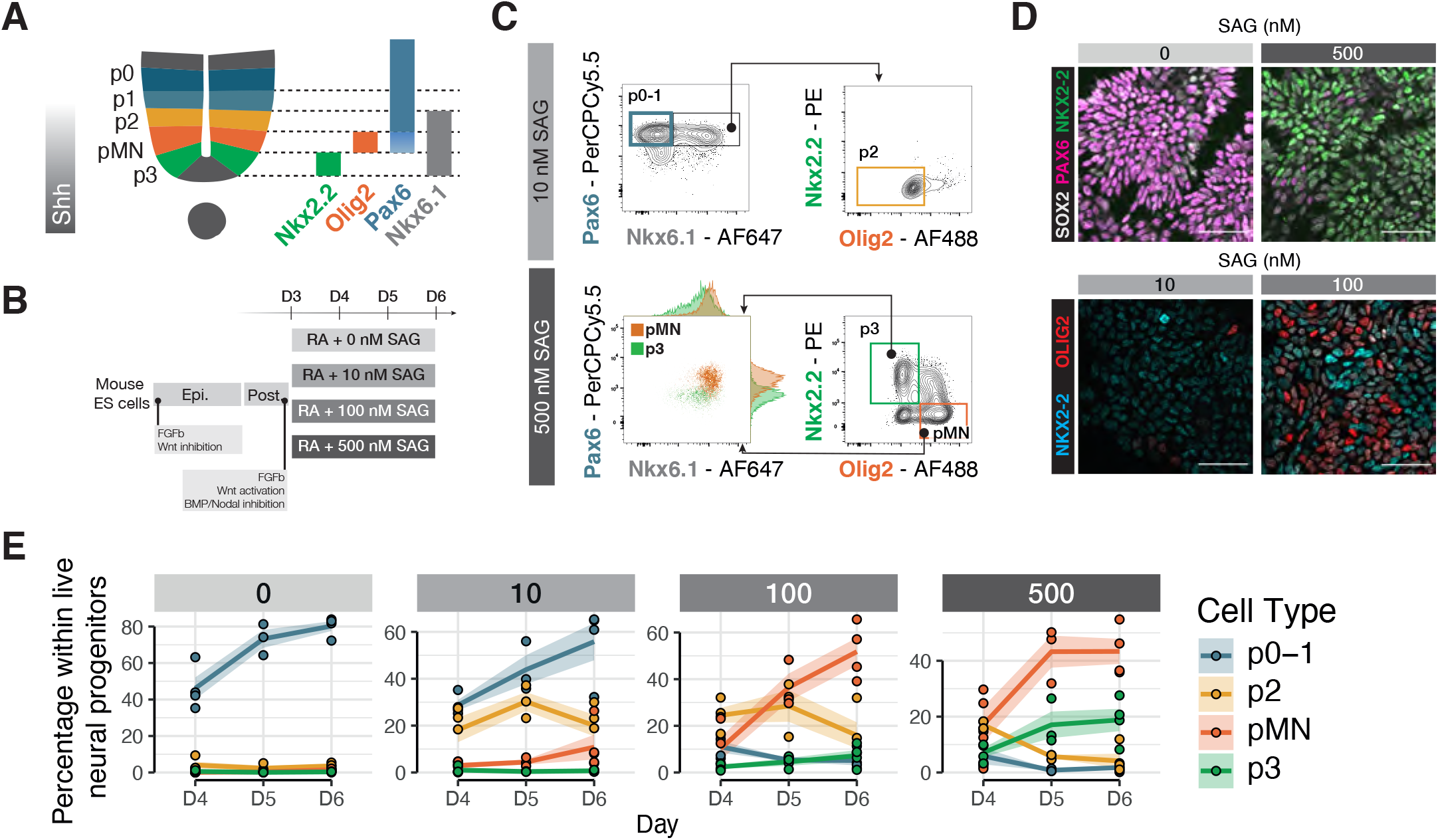
A stem cell model of ventral neural tube progenitors. (A) Schematic of ventral spinal cord progenitors and the markers used for the combinatorial multi-colour flow cytometry and sorting strategies. (B) Schematic of the protocol for the differentiation of mouse ES cells to generate ventral neural progenitors following equivalent signaling cues to embryonic development. (C) Representative flow cytometry plots of the gating strategies used for both analysis and sorting of NPs p0-1, p2, pM and p3. Cells were gated as NPs by excluding dead cells and selecting SOX2+. (D) Representative immunohistochemistry of ES cells differentiated for 6 days show expression PAX6 when exposed to 0 nM SAG and NKX2.2 if exposed to 500 nM. At 100 nM SAG, both OLIG2 or NKX2.2 are detected compared to little or no signal at 10 nM SAG. (E) Proportion of NPs at each SAG concentration shows generation of higher proportions of more ventral cell types at increasing SAG concentrations. Dots are individual samples. Line represents the average. Shaded areas, s.e.m. SAG, Smoothened agonist.

The pattern of progenitor gene expression is determined by a gene regulatory network (GRN). Broadly expressed tissue-wide activators, such as SOX2, promote the transcriptional programmes of multiple progenitor domains (Graham et al., 2003; Oosterveen et al., 2012; Peterson et al., 2012). Concurrently, a set of transcription factors encoding Groucho/TLE-dependent transcriptional repressors are differentially expressed in distinct progenitor domains (Muhr et al., 2001). Pairs of these TFs, expressed in adjacent domains, cross-repress each other and form a network of transcriptional repression that selects a single progenitor identity by repressing all other cell fates (Dessaud et al., 2008; Kutejova et al., 2016; Nishi et al., 2015). The transcriptional effectors of Shh signaling, the GLI (Gli1, -2, -3) proteins provide the spatially polarised input that initiates the patterning process (Hui and Angers, 2011; Persson et al., 2002; Stamataki et al., 2005). The combination of the positive and negative inputs generated by this network establish and position the discrete boundaries of gene expression domains in response to graded Shh signaling (Balaskas et al., 2012; Briscoe et al., 2000; Cohen et al., 2014).

Although the genetic architecture of the GRN is well established, how it is implemented through CREs is less clear. Enhancer elements that drive expression of domain-specific TFs have been identified (Oosterveen et al., 2012; Peterson et al., 2012). The presence of these cell-type-specific enhancer elements could support the Differential Accessiblity strategy in which different CREs are available in different cell types. However, many of these CREs are bound by SOX2, the Shh effector GLI1 and the repressive TFs expressed in alternative cell types, including NKX2.2, OLIG2 and NKX6.1 (Kutejova et al., 2016; Nishi et al., 2015; Peterson et al., 2012). This suggests that the same CREs are employed in different cell types and the composition of TFs at these CREs determines activity: the Differential Binding strategy.

To distinguish between these two regulatory control strategies, we made use of neural progenitors (NPs) differentiated from embryonic stem (ES) cells (Gouti et al., 2014; Sagner et al., 2018). We reasoned chromatin accessibility was the best approach to capture the global landscape of functional regulatory regions. We therefore needed a cell-type-specific accessibility assay, in a system without cell surface markers. Here we developed an ATACseq workflow – Crosslinked and TF-Sorted ATAC-seq (CaTS-ATAC) – based on intracellular flow sorting that allowed the assay of chromatin accessibility of NP cell types. We found that most NPs shared a common set of CREs. These elements are bound by the known repressive TFs, indicating that these cell fate decisions are made by changing the composition of proteins bound to the elements. By contrast, the most ventral NP cell type, p3, has a distinct regulatory programme. We uncover the role of FOXA2 in driving this chromatin re-configuration, and show that it is required for p3 specification. This is reminiscent of the pioneering role of FOXA2 in endoderm lineages, raising the intriguing possibility that it represents a remnant of a GRN co-opted from this germlayer or a shared evolutionary origin for these cell types.

## RESULTS

### A stem cell model of ventral neural tube progenitors

To generate neural progenitors (NP) of different dorsoventral identities we made use of an established protocol for the directed differentiation of mouse ES cells (Gouti et al., 2014; Sagner et al., 2018). To mimic graded Shh signaling we used different concentrations, ranging from 0 nM to 500 nM of the Shh agonist SAG (Fig 1A,B). We observed a dose response as measured by Shh target genes *Gli1* and *Ptch1* (Fig S1A).

Using markers for specific progenitor domains (Delás and Briscoe, 2020), we assayed cell identity with single cell resolution (Fig 1C,D). Each SAG condition generated a mixture of two or three NP types. Using the combinatorial expression of TFs we quantified the proportions of different cell types for each concentration by intracellular staining followed by flow cytometry (Fig 1C). This showed enrichment of the expected cell types. The highest concentration (500 nM SAG) produced the highest percentage of the most ventral domain, p3, characterized by the expression of NKX2.2; pMN, which express the TF OLIG2 were produced at 500 and 100 nM SAG; p2 progenitors, which express NKX6.1 and PAX6, were produced at 100 and 10 nM. PAX6 expression in the absence of NKX6.1 identifies p0 and p1 progenitors and these were produced at 0 and 10 nM SAG. The expected combinatorial expression of TFs was also observed: all p3 and pMN cells also expressed NKX6.1 (Fig 1E). Moreover, we observed a lower level of PAX6 in pMN cells compared to p3 cells, consistent with in vivo data (Fig S1B).

The different SAG conditions also generated the expected neuronal subtypes after addition of Notch inhibitor dibenzazepine (DBZ), which induces neuronal differentiation (Fig S1C). Overall, the data indicate that the ES-derived model of neural tube patterning ventral of progenitors faithfully mimics the in vivo response of neural progenitors to Shh.

### CaTS-ATAC identifies cell type specific regulatory programmes

To distinguish whether the gradient of Shh signaling is interpreted via Differential Accessibility or Differential Binding, we reasoned we needed global chromatin accessibility information with cell type specificity. ATACseq is conventionally performed in live, permeabilised cells. However, the cell type specific markers that distinguish NPs are intracellular and therefore cannot be used on live cells. Based on previous work (Chen et al., 2018) we devised a strategy – Crosslinked and TF-Sorted ATAC-seq (CaTS-ATAC) – to perform ATAC-seq on formaldehyde fixed cells. This involved intracellularly staining and flow cytometry sorting of cells followed by bulk ATAC library preparation (Fig 2A, Methods).

**FIG. 2:**
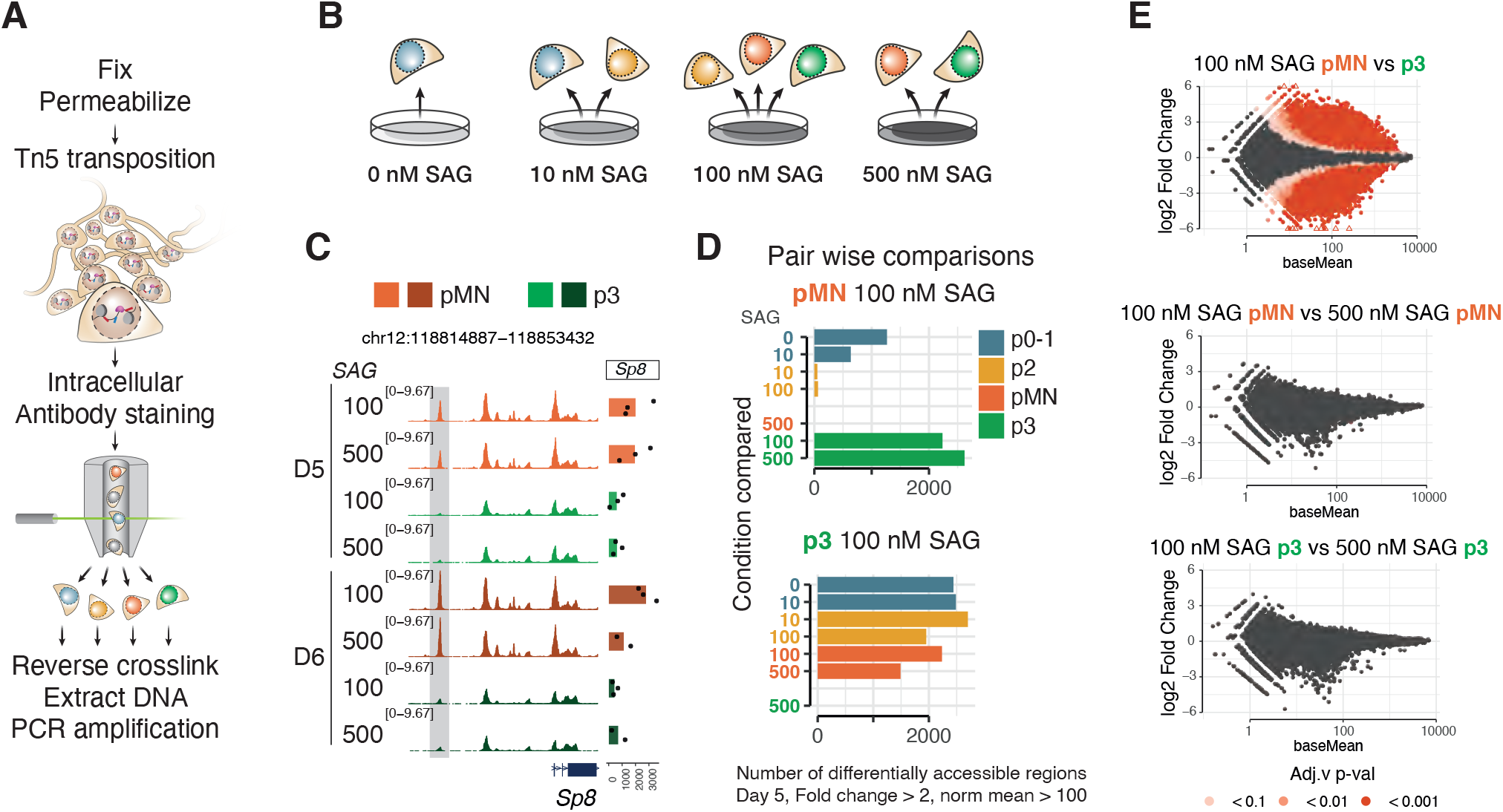
Chromatin accessibility reflects cell type identity independent of SAG concentration. (A) Schematic of CaTS-ATAC, a strategy for cell type specific ATAC-seq based on intracellular markers developed for this study. (B) Cell types analysed from each SAG concentration. (C) Representative genome coverage plot of a differentially accessible region and expression of the nearby gene, Sp8, shows accessibility is consistent for each cell type regardless of the SAG concentration from which it was generated. (D) Quantification of differentially accessible regions between the indicated sample and all other samples at day 5 of differentiation shows no significant differences with the same cell type generated from a different SAG concentration. Thresholds used: absolute fold change > 2, basemean > 100. (E) MA-plot (log2 fold change versus base mean) for the indicated comparisons show large numbers of differentially accessible elements between different cell types generated under the same SAG concentration, but not between the same cell type generated in different SAG concentrations.

We performed CaTS-ATAC-seq over a 4-day timecourse following the addition of different concentrations of SAG. For each condition, we sorted the two or three predominant cell types. This allows us to compare the regulatory landscape of the same cell type originating at different SAG concentrations as well as different cell types within the same dish (Fig 2B).

To explore this dataset, we first examined the correlation between samples based on accessibility at transcriptional start sites (TSS) or distal elements (not overlapping with TSS). As previously reported in other systems (Corces et al., 2016), accessibility of distal elements exhibited greater cell type specificity and higher dynamic range than promoter elements (Fig S2A,B). This confirms our previous observation that ATACseq can capture the regulatory landscape of NPs (Metzis et al., 2018), and it can do so with cell type specificity.

### Cell type specific chromatin configuration is independent of Shh agonist concentration

We first investigated the effect of SAG concentration on cell type specific chromatin accessibility. To do this, we compared global accessibility changes between different NP types generated under the same SAG concentration and the same NP type generated under different SAG concentrations.

We found that the accessibility profiles between the same NP type isolated from different SAG concentrations were indistinguishable (Fig 2C-E). Differential accessibility analysis using DESeq2 revealed no significant differentially accessible peaks between pMN derived from different SAG concentrations (Fig 2D,E), nor between p3 derived from different SAG concentrations (Fig 2D,E). By contrast, over 2000 differentially accessible peaks were identified between pMN and p3 generated under the same SAG concentration (thresholds: fold change over 2, FDR < 0.01 and base mean > 100 normalized counts) (Fig 2D). Similarly, p0-1 or p2 arising from different concentrations had few if any differentially accessible elements (Fig S2C, arrowhead) whereas almost 500 elements were highly differentially accessible between these two NP types generated from the same SAG concentration (Fig S2C).

Each cell type, identified by marker gene expression, acquired a specific chromatin landscape, irrespective of the agonist concentration to which they were exposed. This indicates the GRN mechanism of morphogen interpretation that converts graded Shh input into distinct cell identities establishes not only the discrete transcriptional identities of NPs but also their chromatin landscapes. Thus the differential chromatin accessibility induced by different levels of Shh signaling reflect cell type specific identity and are a product of the GRN that patterns progenitors.

### A shared chromatin landscape for p0-1, p2 and pMN

To examine the global accessibility dynamics, we performed principal component analysis (top 30,000 elements, Fig 3A) and clustered differentially expressed elements across all conditions using k-means (Fig 3B) (Methods). The main change across all data is the global decommissioning of the neuromesodermal progenitor (NMP) programme and establishment of the NP program, which occurs between day 3 and day 4 of differentiation (Fig 3A,B). This involves closing a large proportion of elements open in D3 NMPs (Fig 3B, Cluster 1 “NMP”), as well as opening elements in NPs (Fig 3B, Cluster 3 “Day 4”, Cluster 2 “Day 4-5”, Cluster 9 “PanNeural”). This is expected as cells adopt neural identity. Consistent with this, SOX2 ChIP-seq from NMPs (Blassberg et al., 2022) is enriched in Cluster 1 “NMP” elements and SOX2 ChIPseq from neural cells (Peterson et al., 2012) is enriched in neural clusters (9, 6, 7, 8, 4 and 5) (Fig 3C).

**FIG. 3:**
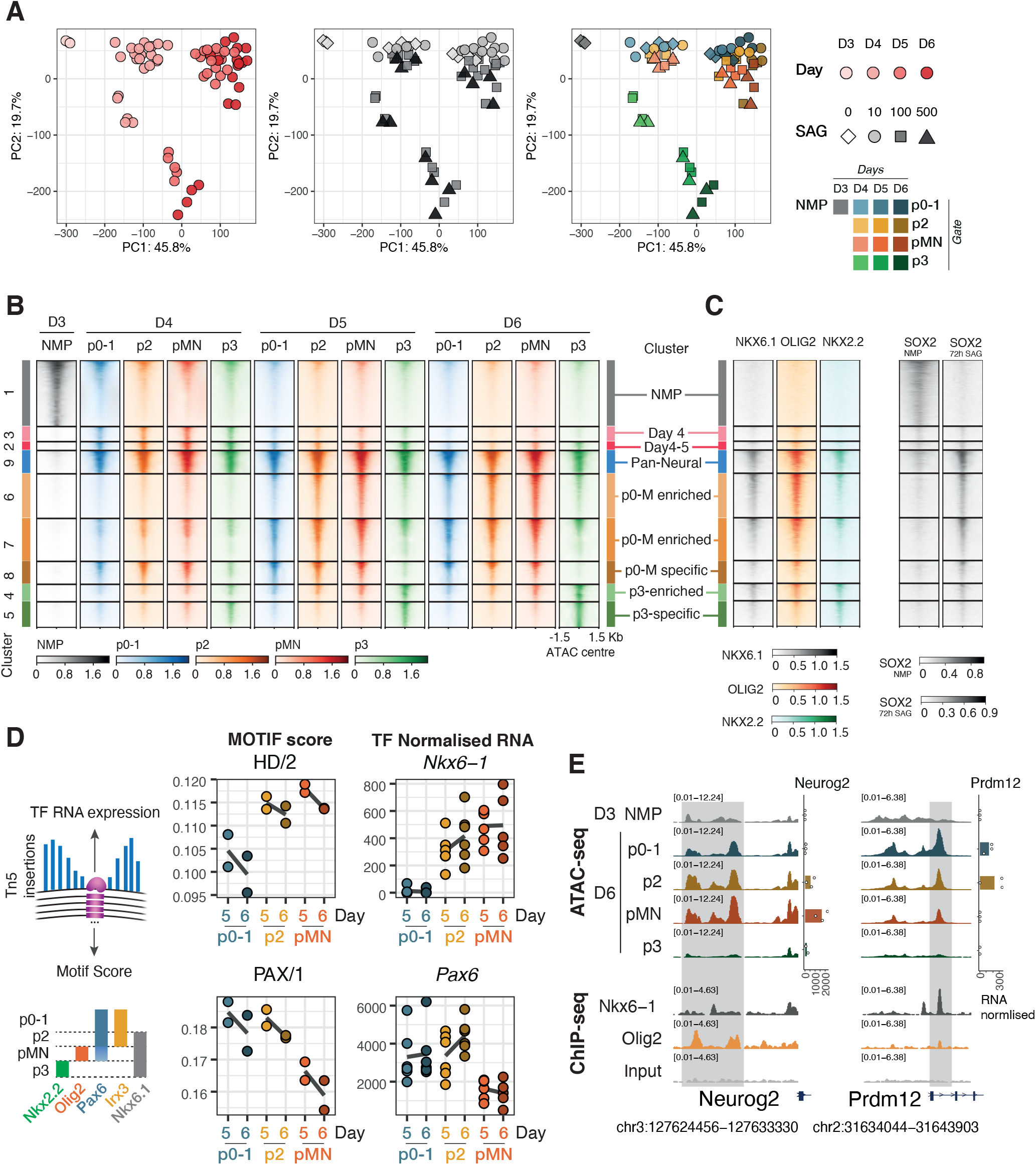
Two regulatory landscapes underlie the Shh response of neural progenitors. (A) Dimensionality reduction (principal component analysis) based on the most variable 30000 consensus elements shows two different behaviours: a shared one for p0-1, p2, pMN samples, and a different one for p3 samples, regardless of SAG concentration. Each symbol represents a sample coloured by differentiation day, SAG concentration or cell type as indicated in the legend. (B) Heatmap showing ATAC-seq coverage for elements differentially accessible between any two cell types or timepoints with the same dynamics, grouped in clusters (see Methods) shows decommissioning of the NMP program (cluster 1), a shared pan-neuronal cluster (cluster 4) and two behaviours in cell type specific accessibility, a shared programme for p0-1, p2 and pMN samples (clusters 6-8), and a unique programme for p3 (cluster 4 and 5). Elements ordered by average accessibility. (C) Heatmap of ChIP-seq coverage for the same elements for NKX6.1, OLIG2, NKX2.2 (Nishi et al., 2015) performed in neural embryoid bodies treated with SAG shows binding to both pan-neuronal and cell type specific elements, and SOX2 from either NMP stage (Blassberg et al., 2022) or SAG treated neural embryoid bodies (Peterson et al., 2012) correlates with accessibility in either NMP or neural progenitors, respectively. (D) Footprinting scores (using TOBIAS, see Methods) for motifs with highly variable scores between p0-1, p2 and pMN at days 5 and 6 obtained, and normalized RNA counts for the most correlated TF within the motif archetype. The motifs for cell type specific TFs show expected footprinting differences.

From day 4 onwards, as cells adopt one of the ventral NP fates, the analysis revealed that pMN, p2 and p0-1 were remarkably similar (Fig 3A,B). This is surprising as these progenitors are molecularly distinct and give rise to functional different neurons: pMN are motor neuron progenitors, characterized by the expression of OLIG2 and generate motor neurons, whereas the p2 domain expresses IRX3 and generates V2 interneurons. RNA expression of marker genes from paired RNAseq samples we also generated confirmed the expected identity of these cell types (Fig S3A). NP p3 were, however, distinct in their chromatin accessibility profile (Fig 3A,B).

The markers that define NP domains are repressive TFs and are directly involved in cell type specification. OLIG2 is expressed in the pMN and represses p3 and p2 fate. NKX6.1 is expressed in p2, pMN and p3 and represses p0-p1. Binding data (ChIP-seq) from these TFs (Nishi et al., 2015) showed both NKX6.1 and OLIG2 bind regions of open chromatin accessible across all the Pan-Neural (Cluster 9) and regions shared by pMN, p2 and p0-1 (p0-M enriched /specific Clusters 6,7,8) (Fig 3C, Fig S3B,C). Since these TFs are only expressed in a subset of the cell types, this suggests that the same elements are open across multiple cell types and occupied by different TFs in different neural progenitor subtypes.

To explore gene regulation differences between p0-p1, p2 and pMN NPs we turned to footprinting. This computational approach uses transposase insertion sites to identify motifs that are under-transposed within open chromatin and are thus likely to be protected by a bound protein. We used TOBIAS (Bentsen et al., 2020) with motifs from three databases (Jolma et al., 2013; Khan et al., 2017; Kulakovskiy et al., 2017), grouping motifs into published archetypes based on position weight matrix clustering of motifs (Vierstra et al., 2020). The analysis revealed homeodomain (HD) and PAX motifs amongst those most differentially scoring between pMN, p2 and p0-1 at day 5 and day 6 (Fig 3D). From all the TFs associated with these motifs, the genes *Nkx6-1* and *Pax6* had RNA expression levels most consistent with the footprint score for HD and PAX, respectively (Fig 3D). In addition, we found the Ebox motif to be amongst the most variable footprints between these NPs (Fig S3D). The high footprint score in pMN is expected as these NP express *Olig2*. But this is not the only Ebox binding TF expressed in these cell types that could explain this footprint. *Neurog2* and *Neurod1* both bind the same motif and are expressed both in p2 and pMN, potentially explaining the Ebox footprint score in p2 NPs. This highlights the challenges in assigning footprints to specific TFs. Overall, the differential footprints found between p0-1, p2 and pMN supports the Differential Binding strategy, in which a common set of open chromatin regions is shared across cell types and different protein composition of the CREs determine gene activity (Fig 3E).

### FOXA2 establishes the p3 regulatory landscape

By contrast to the other NPs, p3 cells had a distinct global accessibility profile, consistent with the Differential Accessiblity strategy (Fig 3A,B). Accessibility was apparent at a unique set of open chromatin regions not shared with any other NP (“p3-specific” Cluster 5), and at a set of elements highly enriched in this cell type (“p3-enriched” Cluster 4). These differences are evident in the second principal component of the PCA where p3 cells are clearly distinct from all other ventral progenitors (Fig 3A).

Analysis of ChIP-seq for the p3-specific TFs NKX2.2 revealed NKX2.2 was enriched in p3-enriched (cluster 4) and p3-specific (cluster 5) (Fig 3C, Fig S3B). However, NKX2.2 was not specific and it also bound the elements shared across all cell types (“Pan-Neural” Cluster 9) and, to a lesser extent, the p0-M-enriches clusters (Clusters 6,7) (Fig 3C, Fig S3B,C). Thus the expression of NKX2.2 did not appear to explain the distinct chromatin landscape of p3. What then drives this distinct chromatin remodelling?

To explore which TFs made p3 NP different from other NPs, we compared footprinting scores across all conditions. The strongest footprint distinguishing p3 from other NPs was predicted to correspond to FOXA2 (Fig 4A,B, Fig S4C). This was intriguing as FOXA2 is known to be expressed during floor plate (FP) development (Altaba et al., 1993; Ang et al., 1993; Sasaki and Hogan, 1993). FOXA2 responds to Shh (Sasaki et al., 1997) and it induces Shh expression (Epstein et al., 1999), thus generating a new signalling centre in the ventral neural tube. To exclude the possibility that the cells isolated as p3 NPs (SOX2+ NKX2.2+ OLIG2-) contained FP cells, we devised culture conditions that promoted expression of mature FP markers (*Arx, Shh*) in which *Foxa2* is expressed at high levels. We compared our spinal cord (SC) differentiation with FP differentiation and observed little expression of FP markers in SC conditions even at the last timepoint sorted, D6 (Fig S4A). Moreover, the expression of FP markers even at later timepoints was very inefficient in SC conditions suggesting that the FOXA2 footprint is not the result of FP induction (Fig S4A,B). We next analysed published FOXA2 ChIPseq generated from neuralised embryoid bodies treated with SAG (Peterson et al., 2012). This revealed FOXA2 binding only to p3-specific peaks (Fig S4D). Consequently, we observed open chromatin in p3 NP samples in the regions bound by FOXA2 (Fig 4C). This contrasts with ChIP-seq of other NP TFs, which showed accessibility across all NP cell types. (Fig S3C). Together, the data prompts the hypothesis that FOXA2 is driving the unique chromatin accessibility profile we observe in p3 cells.

**FIG. 4:**
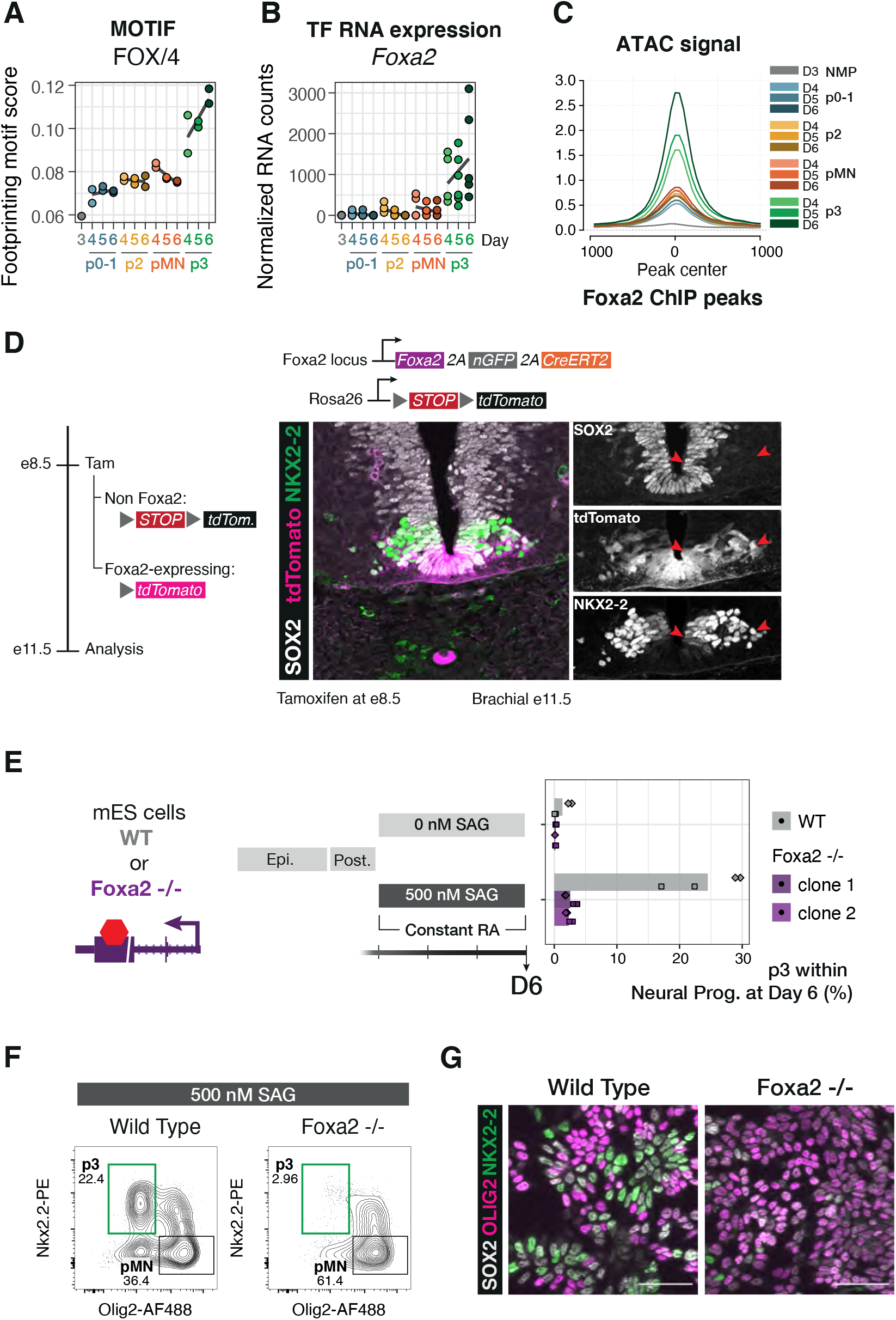
FOXA2 drives the p3-specific chromatin accessibility programme. (A) Footprinting score for the FOX motif is highest in p3 samples. (B) Foxa2 expression in p3 NPs suggests it is the most likely candidate to drive the footprinting signal. (C) Average ATAC-seq accessibility at FOXA2 ChIP-seq peaks (Peterson et al., 2012) in the indicated samples shows these regions are highly accessible in p3 NPs. (D) Genetic lineage tracing indicates that cells that expressed *Foxa2* at E8.5 (tamoxifen administration) have contributed to the p3 progenitor and V3 neuronal cell types by E11.5 (red arrows). (E) *Foxa2*-/- ES cells fail to generate p3 NPs when exposed to 500 nM SAG. (F) Representative flow cytometry plots of the quantifications in (E) showing a marked reduction in p3 NPs from *Foxa2*-/- ES cells compared to wild-type. Cells are gated for SOX2+ live neural progenitors. (G) Representative immunohistochemistry staining for showing reduced number of cells expressing NKX2.2 in *Foxa2*-/- mutant cells at Day 6 of differentiation treated with 500 nM SAG.

### In vivo p3 cells have a history of FOXA2 expression

If FOXA2 plays a role in generating the p3 chromatin accessibility profile, it should be expressed in p3 cells. In vivo NKX2.2 and FOXA2 are co-expressed at early stages of neural tube development (Lek et al., 2010; Ribes et al., 2010) but these FOXA2 expressing cells have been thought to mark the future floor plate. To determine whether cells that express FOXA2 also contribute to the p3 domain, we took advantage of genetic lineage tracing in mouse embryos. Foxa2-driven CreERT2 expression (Imuta et al., 2013) induced recombination of a fluorescent reporter when tamoxifen was administered to pregnant mice at E8.5 or E9.5. Embryos analysed at E11.5 showed expression of the reporter in both p3 (as identified by SOX2 and NKX2.2 expression) and V3 neurons (as identified laterally located cells expressing NKX2.2 alone) (Fig 4D). This demonstrates that p3 cells and their neuronal progeny express *Foxa2* during their history, and supports the hypothesis that FOXA2 expression establishes a unique chromatin landscape in p3.

### FOXA2 is required for p3 cell fate specification

Our results are consistent with the hypothesis that FOXA2 drives extensive remodelling of the chromatin landscape in p3 cells, which in turn is essential for p3 identity. To test this hypothesis, we used genome engineering to create a FOXA2 knockout mouse ES cell line. *Foxa2*-/- cells showed a marked reduction in the proportion of p3 cells (25 % in wild-type to 2 % in the mutants) under 500 nM SAG (Fig 4E-G).

Cells co-expressing NKX2.2 and OLIG2 were still present in *Foxa2*-/- (Fig 4F). This indicates FOXA2 is not directly required for the expression of NKX2.2. Instead, we hypothesise that FOXA2-driven remodelling is required in order for NKX2.2 to exert its repressive activity on OLIG2 and establish the p3 domain. Consistent with this, we observed binding of FOXA2 at the same locations as NKX2.2 (Fig S4E).

### FOXA2 expression substitutes for Shh signalling early in p3 specification

FOXA2 is expressed in NPs closest to the source of Shh and it is only co-expressed with NKX2.2 at early stages of neural tube development. To ask whether early exposure to Shh signaling is necessary to establish a p3 identity, we took advantage of our in vitro NP differentiation protocol to investigate the timing requirements for FOXA2 expression and Shh signaling in p3 specification. To this end, we exposed cells to 0 nM SAG for 24h before changing regime to 500 nM SAG. Compared to a constant exposure to 500 nM SAG, cells exposed to delayed SAG are greatly impaired in their generation of p3 NPs (Fig S5A,B). This was not due to reduced Shh transduction (Fig S5B). The reduction in p3 specification was reminiscent of the outcome of exposing *Foxa2*-/- NPs to 500nM SAG (Fig 4E,F) as we also saw NPs co-expressing NKX2.2 and OLIG2 (Fig S5B). When cells were exposed to SAG from day 3, a proportion of them show low but detectable levels of FOXA2 after 24h. In the delayed SAG addition conditions, however, cells did not upregulate FOXA2 in the equivalent timeframe (Fig S5D).

This regime provided the opportunity to test whether FOXA2 expression within the initial 24h of NP differentiation was sufficient to rescue p3 generation. We used doxycycline inducible lines that expressed either mCherry or a Foxa2-mCherry fusion under the control of the doxycycline-responsive promoter tetON. Consistent with prior experiments, overexpression of mCherry alone in this “delayed SAG” regime showed a reduction in p3 generation at D6 (Fig 5A). By contrast, enforced expression of *Foxa2* for 12h during the initial 24h of neural differentiation rescued p3 generation to levels comparable to those observed with constant 500 nM SAG (Fig 5A).

**FIG. 5:**
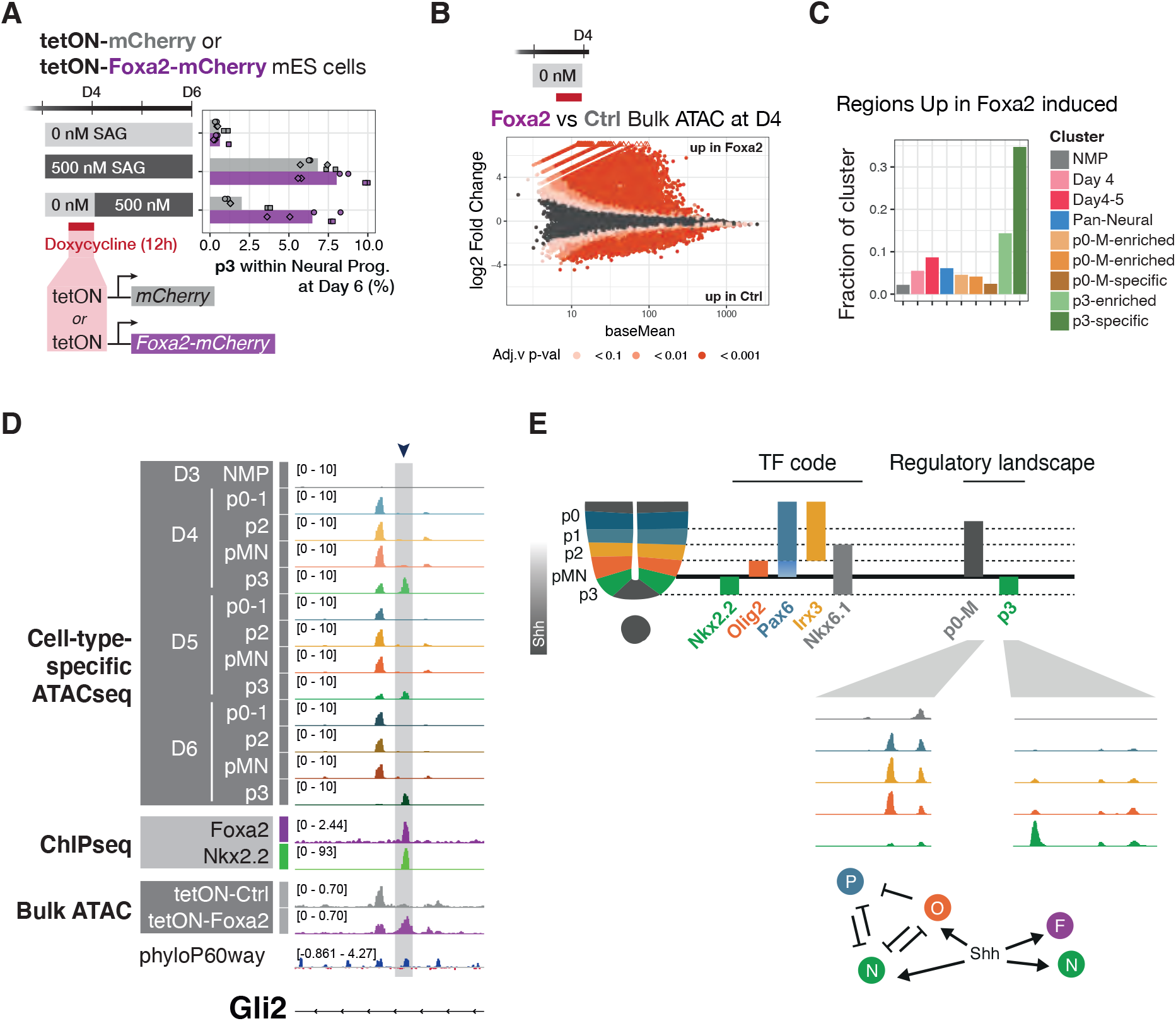
FOXA2 can replace Shh in early p3 specification where it opens p3-specific regulatory elements. (A) A delayed SAG regime greatly reduces p3 generation in control cells (mCherry). A 12h overexpression of tetON-Foxa2-mCherry rescues p3 generation. (B) Differential accessibility in Foxa2 forced expression versus control mCherry shows FOXA2 predominantly opens elements. (C) Overlap of differentially upregulated regions from (B) with ATAC clusters (Fig 3B) reveals a large fraction of the p3-specific cluster is opened upon Foxa2 overexpression. (D) Example element in a *Gli2* intron with p3-specific chromatin accessibility, FOXA2 and NKX2.2 binding that gains accessibility upon *Foxa2* overexpression but not control. (E) Diagram describing the two regulatory landscapes that underlie the molecular identity of ventral neural progenitors, a differential binding strategy is used to distinguish p0-1, p2, pMN, whereas p3 employ a differential accessibility strategy.

These data indicate that high Shh at the onset of spinal cord progenitor specification is required to induce *Foxa2*. This expression of FOXA2 is sufficient to confer competence for Shh signaling to specify p3 identity.

### FOXA2 is sufficient to remodel the chromatin landscape

To test whether FOXA2 is responsible for the early p3-specific chromatin remodelling, we took advantage of the tetON cell lines. We performed bulk ATACseq at D4, which is 24h after the induction of neural identity and 12h after doxycycline exposure (Fig 5B). Both control and FOXA2-overexpressing conditions showed accessibility across the previously defined NMP and pan-neuronal clusters (Fig S5F). Although neither cell line had been exposed to SAG, FOXA2-expressing cells displayed opened chromatin across the p3-specific cluster (Fig S5F).

Consistent with its known role as a pioneer factor (Cirillo et al., 2002), FOXA2 overexpression resulted in chromatin opening (Fig 5B). Over 30% of the regions in the p3-specific cluster overlapped with regions more accessible in FOXA2-overexpressing cells (Fig 5C). These results support a model in which FOXA2 opens chromatin regions required for the p3 fate. Thus Shh induction of *Foxa2* at early neural developmental stages is required to configure the p3-specific response of the cells to Shh.

### A common lineage-pioneering role for FOXA2 across germ layers

FOXA TFs are required for the development of endoderm-derived lineages, pancreas, liver and lung, where they play partially redundant roles (Cernilogar et al., 2019; Geusz et al., 2021; Lee et al., 2005, 2019; Wan et al., 2005; Wang et al., 2015). Whereas many TFs play roles in multiple tissues, they are generally thought to act via different enhancers in a combinatorial fashion with other tissue-specific TFs (Meredith et al., 2013; Zinzen et al., 2009). Pioneer TFs are proposed to act by opening up compacted chromatin, thus allowing other TFs to bind (Zaret and Carroll, 2011). Thus, a given pioneer TF could have access to the same binding sites independent of the cellular context. However the binding of pioneer TFs has been shown to depend on epigenetic features (Lupien et al., 2008). Indeed cell type specific co-factors direct cell-type-specific binding of FOXA2 in endogenously-expressing cell lines and in FOXA2 overexpressing cell lines (Donaghey et al., 2018).

We therefore asked how much of FOXA2’s pioneering activity is tissue specific between endoderm and neural p3 progenitors. We examined the binding of FOXA2 in cellular differentiation models of endoderm at day 3 and day 5 (Cernilogar et al., 2019), and asked whether endoderm FOXA2 was bound to any of the spinal cord differentiation clusters. To our surprise, we found FOXA2 also bound to the p3-specific cluster of elements in endoderm (cluster 5) (Fig 6A). This included genes such as *Prox1*, expressed in both lineages (Burke and Oliver, 2002; Kaltezioti et al., 2014); *Lmx1b*, required for specification of hindbrain p3-derived serotonergic neurons (Cheng et al., 2003; Ding et al., 2003), and for in vitro pancreatic islet generation (Alvarez-Dominguez et al., 2020); and *Rfx3*, also required for pancreatic endocrine differentiation (Ait-Lounis et al., 2007) and expressed in the ventral neural tube (Cruz et al., 2010) (Fig 6B). This argues for a functional role in both tissues for at least some of these targets. Moreover, many of these sites were either opened in neural progenitors upon brief over-expression of *Foxa2* (Fig 6B, Fig S5F), or occupied by overexpressed FOXA2 in endoderm differentiation (Fig 6B).

**FIG. 6:**
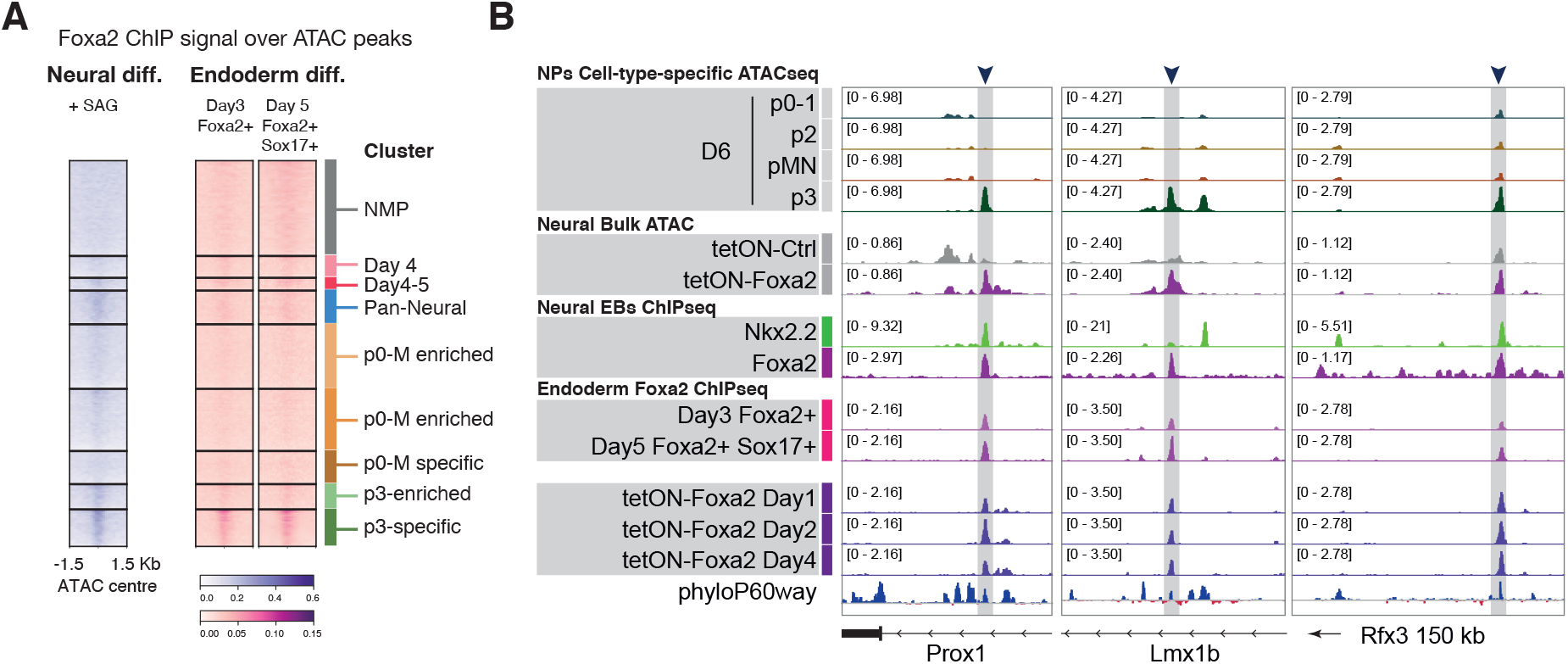
A common regulatory role of FOXA2 in ventral neural and endoderm lineages. (A) Heatmap of ChIP-seq coverage for neural FOXA2 and endoderm FOXA2 at two differentiation timepoints (Cernilogar et al., 2019) show binding for both in the p3-specific accessibility cluster. (B) Potentially functionally relevant target genes expressed and/or required in both tissues, with p3-specific accessibility, opened in response to FOXA2 overexpression, bound by FOXA2 in neural and endoderm, including overexpression of FOXA2.

While endoderm and neural progenitors also have tissue-specific FOXA2 binding, these data raise the possibility of the two tissues share a core set of targets. Mechanistically, this could be mediated by the classical pioneer role of FOXA2 in this subset of targets, or by the presence of common co-factors in both tissues. Intringuingly, it could offer evidence of an evolutionarily conserved GRN.

## DISCUSSION

Allocating cell fates in a developing tissue involves the ordered specification of multiple, alternative cell identities. A well characterised example is neuronal subtype generation in the ventral neural tube, directed by the morphogen Shh. Here we uncover two cis regulatory strategies by which graded Shh signaling directs cell fate choice in neural cells. One strategy – Differential Binding – relies on a common regulatory landscape. The different composition of TFs at these CREs dictates differential gene expression and cell fate decisions. We show that neural progenitors p0-1, p2 and pMN use this strategy, acting through a set of shared accessible chromatin regions bound by cell type specific repressive TFs. The second strategy – Differential Accessiblity – relies on cell type specific chromatin remodelling. This is the case for p3 NPs, which have a unique set of accessible elements that distinguish them from all other NPs. We show that FOXA2, expressed early in response to Shh, is responsible for the remodelling that opens p3-specific elements. Many of these elements then bind the p3-specific TF, NKX2.2. Intriguingly, a subset of the elements bound and opened by FOXA2 in neural progenitors are also bound by FOXA2 during endoderm differentiation, where it plays a parallel role opening elements later required for specific endoderm lineages (Geusz et al., 2021).

To assay cell type specific chromatin accessiblity we took advantage of the combinatorial TF code that controls ventral spinal cord specification. Since TFs are intracellular, this required developing methods to perform ATAC-seq from paraformaldehyde-fixed cells, followed by intracellular flow cytometry with a set of six antibodies that allowed purification of specific NP subtypes using the combination of TFs they express. This approach is based on previous methods developed to measure protein levels at single cell resolution using plate-based assays (Chen et al., 2018) and produced high quality chromatin accessiblity data for specific cell types (Baskar et al., 2022). The methods developed here are applicable to other tissues, in vitro and in vivo, and will be of particular use in cases where fluorescent reporters are not readily available or when the identification of the desired cell types depends on the intersectional expression of multiple markers. Moreover, the approach could be extended to isolate cells based on other intracellular characteristics, for example post-translationally modified signalling effectors.

A common set of pan-neural elements, not accessible prior to neural induction, was apparent in all NPs, irrespective of their identity. These elements were enriched for binding of TFs involved in the neural tube GRN, including SOX2, NKX6.1, NKX2.2 and OLIG2. The existence of this set of elements is reminiscent of the idea of morphogenetic fields – discrete, modular units of embryonic development in which signaling coordinates the spatial organisation of cells (Gilbert et al., 1996). A common chromatin landscape could provide a molecular correlate of a morphogenetic field and explain the coordinated response of a field of cells to patterning signals. Moreover, comparison of chromatin accessiblity in the same NP cell type generated using different levels of Shh signaling revealed that cell identity, rather than morphogen concentration, correlated with chromatin accessiblity pattern. This indicates that the morphogen interpretation mechanism that converts graded Shh input into distinct cell identities establishes not only the discrete transcriptional identities of NPs but also their chromatin landscapes.

The majority of ventral NPs shared the same overall chromatin landscape, suggesting a Differential Binding strategy governs gene expression selection in these cases. This is consistent with the ‘selection by exclusion’ mechanism that has been proposed previously to explain the specification of progenitor subtype identity by graded Shh signaling (Kutejova et al., 2016; Nishi et al., 2015). In this view, the positive and negative transcriptional inputs supplied by the GRN are integrated at CREs associated with target genes, which are accessible in NPs regardless of whether the target gene is active or not. Thus, cell fate choice does not depend on major chromatin remodelling, but instead is determined by the composition of transcriptional effectors bound.

By contrast, the Differential Accessibility strategy appears to dictate the choice of p3 NPs. These NPs are distinguished by accessiblity at a distinct subset of CREs that depend on FOXA2. Despite the transience of FOXA2 expression in p3 NPs (Wang et al., 2011), accessibility at the CREs opened by FOXA2 persists and are bound by the p3-specific TF NKX2.2. This is similar to the observation that the presence of FOXA is not required to maintain stable epigenetic states in liver cells (Reizel et al., 2021). The marked changes in chromatin landscape initiated by FOXA2 thus represent an example of epigenetic memory and demonstrate how the pioneering activity of a TF can rewire a GRN to specifiy a specific cell type. A consequence of this is that cell fate is not simply the product of the TFs it expresses, but also the result of its gene expression history.

The activity of FOXA2 may also account for the different neuronal subtypes generated from p3 NPs. Although in the spinal cord p3 NPs produce local projecting excitatory interneurons that are part of the motor control circuitry (Zhang et al., 2008), in the hindbrain p3 NPs generate serotonergic neurons (Briscoe et al., 1999), and midbrain FOXA2-expressing NPs generate dopaminergic neurons (Kittappa et al., 2007). The capacity of FOXA2 to modulate chromatin accessiblity and thereby rewire the GRN may explain the diversity of neuronal subtypes produced by p3 NPs along the rostral caudal axis.

Two implications for cellular reprogramming emerge as a consequence of the different regulatory strategies. Cell types sharing the same landscape (Differential Binding) would be relatively plastic, enabling transitions between the different fates by altering the TF configuration on already available CREs. Consequently, the barrier to reprogram cells between such fates would be lower. On the other hand, the Differential Accessibility strategy involves chromatin remodelling to facilitate the binding of TFs previously unable to bind specific CREs. The expression of the end point TFs alone might not be sufficient for cell fate acquisition if the elements these TFs need to bind are inaccessible. This might explain the inefficiency or incompleteness of some transgene-driven reprogramming approaches that use only the TFs expressed in the final state of the target cell type (Nashun et al., 2015). Understanding and recapitulating the epigenetic trajectory cells follow to reach the desired endpoint might improve such differentiation protocols.

Our results are consistent with prior genetic studies showing the p3 NPs are the only NPs that require activating GLI proteins for their specification (Litingtung and Chiang, 2000; Persson et al., 2002). Embryos lacking Shh and the main source of repressive GLI activity, GLI3, are able to generate all ventral NP types, except p3 (Litingtung and Chiang, 2000; Persson et al., 2002). The demonstration that FOXA2 expression is able to substitute for Shh signaling during the early steps of p3 specification indicates that the requirement for high levels of Shh signaling is to initiate FOXA2 expression. This requirement restricts p3 induction to the most ventral progenitors in the neural tube. The expression of FOXA2 and NKX2.2 in p3 NPs also appears to contribute to establishing p3 identity by modulating the cell intrinsic response to Shh (Lek et al., 2010). NKX2.2 promotes *Foxa2* via a mechanism involving *Gli3* repression (Lek et al., 2010) and FOXA2 binds and opens an intronic region in the gene encoding *Gli2* (Fig 5D). Together, these data suggest that the interpretation of Shh signaling by p3 NPs is distinct from the adjacent p0-pMN region. This distinction involves remodelling the regulatory landscape of p3 NPs.

FOXA2 has been proposed to act as a pioneer factor in the endoderm, where it remodels chromatin to allow the binding of other TFs (Geusz et al., 2021; Zaret and Carroll, 2011). Although TFs often have roles in multiple tissues, they are usually thought to act in a combinatorial fashion with other TFs via different elements in different tissues (Donaghey et al., 2018; Meredith et al., 2013; Zinzen et al., 2009). The observation that FOXA2 binds to p3 NP specific CREs in endodermal cells raises the possibility these two cell types share some aspects of their regulatory control. Moreover, the role of FOXA2 in endoderm is highly evolutionarily conserved (Stainier, 2002). Pancreas specification, an endoderm derivative, involves many TFs that are also expressed in ventral neural tube, specifically in the p3 domain (Larsen and Grapin-Botton, 2017). The potential shared control by FOXA2 of functionally relevant genes in endoderm and neural tissue raises the intriguing possibility of co-option of an endoderm FOXA2 regulatory programme into neural progenitors. Alternatively, it could be evidence of a common evolutionary origin for endoderm and neural cells. This would be supported by recent work suggesting that parts of the digestive system in the cnidaria Nematostella are of ectodermal origin (Steinmetz et al., 2017), and that a common progenitor generates neurons and secretory cells in this species (Tournière et al., 2022) (Steger et al., 2022). A deeper understanding of the GRNs in neural tissue and endoderm will be needed to explore this possibility. More generally it will be important to determine whether other tissues use similar cis regulatory strategies to specify distinct cell types.

## Supporting information

Oligo_table

## ACKNOWLEDGEMENTS

We are grateful to A. Sagner, V. Metzis and T. Rayon, and members of the Briscoe lab for advice and reagents. We thank Anne Grapin-Botton and Uli Technau for critical reading of the manuscript. We thank the Crick Science Technology Platforms, in particular, Vangelis Christodoulou from the Structural Biology STP for production of Tn5 enzyme, Flow Cytometry, Advanced Sequencing, Advanced Light Microscopy, BRF and Advanced Computing. This work was supported by the Francis Crick Institute, which receives its core funding from Cancer Research UK, the UK Medical Research Council and Wellcome Trust (all under FC001051). Work in the Briscoe lab is funded by the European Research Council under European Union (EU) Horizon 2020 research and innovation program grant 742138. This work was also funded by the Wellcome Trust (220379/D/20/Z). H.T.S. is supported by a Sir Henry Wellcome postdoctoral fellowship.

## AUTHOR CONTRIBUTIONS

M.J.D and J.B. conceived the project, interpreted data and wrote the manuscript. M.J.D. designed and performed experiments, performed bioinformatic analysis and data analysis. C.M.K., T.F., M.D. and I.Z. performed experiments. H.T.S., E.C. and E.M.T. generated and characterized the HM1-tetON cell lines. K.I. designed and assisted with embryology experiments.

## EXPERIMENTAL PROCEDURES

### Experimental Model and Subject Details

#### Lineage tracing of Foxa2-expressing cells

Foxa2-^2a^-^nEGFP-2a-CreERT2/+^;R26^Tomato Ai14/ Tomato Ai14^ embryos were obtained from time matings. Mouse lines Foxa2^nEGFP-CreERT2/+^ (MGI:5490029) (Imuta et al., 2013) and R26^tdTomatoAi14/tdTomatoAi14^ (Gt(ROSA)26Sor^tm14(CAG-tdTomato) Hze^ (MGI:3809524) (Madisen et al., 2010) were maintained in a C57BL6 background. Induction of recombination was achieved by tamoxifen administration by oral gavage at 0.08 mg/body weight.

All animal procedures were performed in accordance with the Animal (Scientific Procedures) Act 1986 under the UK Home Office project licenses PP8527846 and PF59163DB.

#### Cell lines

WT experiments were performed with the mouse embryonic stem cell line HM1 (Doetschman et al., 1987). All cell lines were maintained at 37^*◦*^C with 5% carbon dioxide (CO_2_).

#### Generation of Foxa2^-/-^ ES cell lines

Generation of Foxa2 knockout cell lines was performed by electroporating two different guides cloned into the pX458 vector (Ran et al., 2013) with the AMAXA nucleofector kit (Lonza Cat no. VPH-1001). Cells were sorted for GFP as single cells one day after electroporation. Clonal cell lines were genotyped and two clones were used. Both carried the same deletion.

#### Generation of tetON-mCherry and tetON-Foxa2-mCherry ES cell lines

PiggyBac (PB)-tetOn-destination-PGK-hygro and PB-CAG-rtTA3-PGK-puro vectors were used from (Stuart et al., 2019). Gateway cloning (Invitrogen) was used to insert coding sequence for mCherry or FoxA2:mCherry fusion protein into the tetOn destination vector. Stable transgenic HM1 mouse ES cell lines were generated by transfection as follows: 1 *µ*g PB-tetOn expression vector, 1 *µ*g PB-CAG-rtTA3 vector, 1 *µ*g non-integrating PBase transposase vector and 2 *µ*l Lipofectamine-2000 (Invitrogen) were incubated for 20 min in 100 *µ*l DMEMF12 (Gibco) at room temperature, then applied to 300,000 cells/6well in ml medium for 18 hours. Selection was applied to transfectants for at least 5 passages prior to use: 150 *µ*g/ml hygromycin-B (ThermoFisher) and 1 *µ*g/ml puromycin (ThermoFisher). Transfection and selection were performed in feeder-free 2iLIF culture conditions as described in (Stuart et al., 2019), then adapted back to feeders + ES cell medium regime as described below for at least 4 passages prior to experiments.

### Method Details

#### Cell culture and neural progenitor differentiation

Mouse ES cell lines were maintained in ES cell medium (Dulbecco’s Modified Eagle Medium (DMEM) Knock Out (Gibco, Cat No. 10829-018) complemented with 10% Foetal Bovine Serum (Pan Biotech, Cat. No. P30-2602), Penicillin/Streptomycin (Gibco, Cat No. 15140122), 2mM L-Glutamine (Gibco, Cat No. 25030024), 2mM Non-essential amino acids (Gibco, Cat No. 11140-035), and 0.1 mM 2-mercaptoethanol (Gibco, Cat. No. 21985-023)) with 1000 U/ml LIF (Chemicon, Int ESG1107) on mitotically inactive primary mouse embryo fibroblasts (feeders). For spinal cord neural differentiation, ES cells were dissociated using 0.05% trypsin-EDTA (Gibco, Cat No. 25300054) and then plated onto 0.1% gelatinised (Gibco Cat no. G1393-100ML) tissue culture plates for three successive periods of 15 mins to remove feeders. Differentiations were carried out in N2B27 media (Advanced DMEM - F12 (Gibco, Cat. No. 21331-020) and Neurobasal medium (Gibco, Cat. No. A35829-01) (1:1), supplemented with 1xN2 (Gibco Cat no. 17502001), 1xB27 (Gibco Cat no. 17504001), 2 mM L-glutamine (Gibco, Cat No. 25030024), 40 *µ*g/ml BSA (Sigma-Aldrich, Cat No. A7979-50ML), and 0.1 mM 2-mercaptoethanol) with the indicated additives. Cells were plated onto 6-well plates (Corning, Cat. No. 353046) precoated in Matrigel (Corning, Cat. No. 356231) diluted 1/50 and 1/100 in Advanced DMEM - F12 at a density of 20,000 cells per 35mm well in 1.5 ml. The media was supplemented on the different days as follows: Day 0 to Day 2, 10ng/ml bFGF (R&D, Cat. No. 100-18B) and 5 *µ*M LGK (Cayman Chemical Company, Cat. No. 1.800.364.9897); Day 2 to Day 3 for 20 h, 10 ng/ml bFGF, 5 *µ*M CHIR99021 (Axon Medchem, Cat. No. 1386), 10 *µ*M SB-431542 (Tocris, Cat. No. S0400), and 2*µ*M DMH1 (Adooq Bioscience, Cat. No. A12820); from Day 3 onwards, 100 nM RA (Sigma, Cat. No. R2625) and either 0 nM, 10 nM, 100 nM or 500 nM SAG (Merck, Cat. No. 566660-5mg). The “delayed SAG” regime consisted of 100 nM RA and 0 nM SAG from Day 3 to Day 4, followed by a constant concentration of RA and increased SAG to 500 nM. Induction of the tetON system was achieved by supplementing the media with 1 *µ*g/ml Doxycycline (Sigma-Aldrich, Cat. No. D9891) for the second 12h of Day 4. Floor plate conditions were achieved by addition of 500 nM SAG on Day 3 without any addition of RA.

#### Flow cytometry of intracellular markers

##### Sample collection

1*µ*l/ml of LIVE/DEAD™ Fixable Dead Cell Stain Near-IR fluorescent reactive dye (ThermoScientific, Cat. No. L34976) was added to cells in culture and incubated at 37^*◦*^C for 30 mins. After cells were washed twice with Phosphate Buffer Saline (PBS) (Gibco, Cat. No. 14190-094), cells were dissociated using 0.5ml Accutase (Gibco, Cat. No. 00-4555-56) per well of a 6 well plate incubated 5 min at 37^*◦*^C. Cell were collected, centrifuged at 400g for 4 min and resuspended in 100*µ*l of 4% paraformaldehyde (PFA) (ThermoScientific, Cat. No. 28908). PFA fixation was carried out for 10 min at room temperature. Cells were washed in PBS and resuspended in 500*µ*l PBS + 0.5% BSA.

##### Staining

1 million cells were stained for flow cytometry analysis. Pellets were resuspended in 0.1ml of PBS + 0.5% BSA and 0.1% Triton-X100 (VWR Chemicals, Cat No. 28817.295) and were incubated on ice for 10 mins. Cells were then incubated with primary antibodies or directly conjugated antibodies diluted in the same buffer for 1.5 h protected from light at room temperature. When necessary, secondary antibodies were incubated under the same conditions for 40 min. Cells were washed in PBS + 0.5% BSA and 0.1% Triton-X100, and resuspended in 300*µ*l of PBS + 0.5% BSA for analysis on a BD Fortessa analyzer (Becton Dickinson). The antibody panels used were as follows: “D-V progenitors”, Sox2-V450 (1:100), Pax6-PerCPCy5.5 (1:100), Nkx6.1-AlexaFluor647 (1:100), Goat Olig2 unconjugated (1:400), Nkx2.2-PE (1:100), and Tubb3-Biotin (1:800) followed by secondaries donkey anti-goat AlexaFluor488 (1:1000) and Strep-APC-Cy7 (1:20,000); “mCherry inducible”, Sox2-V450 (1:100), Pax6-PerCPCy5.5 (1:100), Goat Olig2 unconjugated (1:400), Nkx2.2-AlexaFluor647(1:100), secondaries donkey anti-goat AlexaFluor488 (1:1000); “D-V with Foxa2”, Sox2-V450 (1:100), Pax6-PerCPCy5.5 (1:100), Goat Olig2 unconjugated (1:400), Nkx2.2-PE (1:100) and Foxa2-AlexaFluor488 (1:100), with secondary donkey anti-goat AlexaFluor647 (1:1000). Antibody details in Table S1.

#### RNA extraction, cDNA preparation and qPCR analysis

Wells were washed with PBS and lysed in 350 *µ*l of RLT buffer (QIAGEN, Cat. No. 1015762) per 35mm well. After cells were mixed, they were collected in a 2 ml RNase-free Eppendorf tube and stored at -20^*◦*^C. RNA extraction was performed using RNeasy Qiagen kit with DNAse digest (QIAGEN, Cat. No. 74106) as per manufacturer’s protocol. cDNA were synthesized from 1ug of RNA using Superscript II reverse transcriptase (Invitrogen 18080-044) with random hexamers, and qRT-PCR analyses were performed by QuantStudio 12K Flex Real-Time PCR system (ThermoFisher Scientific) using SYBR Green PCR assay (ThermoFisher Scientific, Cat. No. A25742). All experiments were performed in technical duplicates, biological duplicates or triplicates for each time point analysed. Expression values were normalized against *β*-actin. Primer sequences in (Table S2).

#### Immunofluorescence staining of cell differentiations

Cells for immunofluorescence staining were cultured in matrigel-coated glass slides in 12-well plates with all the volumes adjusted to achieve the same cell density. Wells were washed with PBS and fixed in 4% PFA at 4^*◦*^C for exactly 15 mins. Glass slides were transferred to a new 12-well plate for staining. Blocking was performed for 10 mins with PBS + 1% BSA + 0.1% Triton-X100 at room temperature, cells were incubated in PBS + 1% BSA + 0.1% Triton-X100 with primary antibodies overnight at 4^*◦*^C, with secondary antibodies at room temperature for 1hr, and mounted with ProLong Gold antifade reagent (Invitrogen, Cat. No. P36930). Fluorescent images were taken with Leica SP8 confocal microscope or an Apotome microscope.

The following primary antibodies were used: goat anti-Sox2 (1:500), mouse anti-Nkx2.2 (1:1000), rabbit anti-Olig2 (1:500), rabbit anti-Pax6 (1:1000) (4C fridge), Foxa2-AlexaFluor488 directly conjugated (1:100). Corresponding donkey raised secondaries. Antibody details in Table S1.

#### Generation of mouse sections and immunostainings

Mouse spinal cord sections staining were performed as previously described (Rayon et al., 2020). Mouse embryos from timed pregnant females were collected and fixed in 4% PFA for 1.5h at 4^*◦*^C, washed in PBS, and transferred to 15% sucrose in phosphate buffer overnight at 4^*◦*^C. Embryos were subsequently embedded in gelatin solution (7.5% gelatin, 15% sucrose in phosphate buffer) and snap-frozen in isopentane on dry ice. Transverse cryosections (thickness: 14 mm) were cut using a Leica CM3050S cryostat (Leica Microsystems Limited, Milton Keynes, UK) and placed on Superfrost Plus slides (Thermo Scientific, Cat. No. 10149870). Slides were stored at -80^*◦*^C until ready to be processed for immunohistochemistry. After immunohistochemistry, 22-mm– by–50-mm no. 1.5 thickness coverslips (VWR, Cat. No. 631-0138) were mounted onto the sections using ProLong Gold antifade reagent. Immunohistochemistry was performed in the same way as for cell culture slides using the same antibodies against Sox2 and Nkx2.2.

#### CaTS-ATAC (Crosslinked and TF-Sorted ATAC-seq)

##### Sample collection

Cell type specific ATAC-seq based on intracellular markers was carried out from HM1 differentiations at the timepoints and SAG concentrations indicated. Dishes were washed with PBS and dissociated with 0.5ml Accutase (Gibco, Cat. No. 00-4555-56) per well of a 6 well plate incubated 5 min at 37^*◦*^C. Cells were collected in Eppendor LoBind tubes (Eppendorf cat. #Z666548), the wells were rinsed with 1ml PBS and the samples were spun at 400 g for 4 min at room temperature. Samples were resuspended in 1*µ*l/ml of LIVE/DEAD™ Fixable Dead Cell Stain Near-IR fluorescent reactive dye as per manufacturer’s instructions. After 30 min incubation on ice protected from light the samples were spun again and resuspended in 300 *µ*l of PBS, followed by addition of 100 *µ*l of PFA to achieve 1% PFA. Fixation was carried out for 15 min with rotation at 4^*◦*^C. Fixation was quenched with 25 *µ*l of 2M Glycine for 5 min at 4^*◦*^C with rotation, spun and resuspended in PBS with 0.5% BSA. Cells were counted and 1 million cells were transferred to a new LoBind Eppendorf tube.

##### Transposition

Tn5 transposition was carried out as previously described (Corces) with some modifications. Briefly, samples were resuspended in 0.5 ml of RSB buffer (10 mM Tris-HCl pH 7.4, 10 mM NaCl, 3 mM MgCl2) supplemented with 0.1% Igepal CA-630 (Sigma, Cat. No. I8896-100ML), 0.1% Tween-20 (Sigma, Cat. No. P2287-500ML), and 0.01% digitonin (Invitrogen, Cat No. BN2006), and incubated on ice for 3 min. 1 ml of RSB with 0.1% Tween-20 was then added and samples were spun for 10 min at 2000 g and 4^*◦*^C. The samples were then resuspended in 1 ml of ATAC mix (2X TDE buffer (Illumina), 50 *µ*l TDE (Illumina), 0.01% digitonin, 0.1% Tween-20 and 330 *µ*l of PBS). Transposition was carried out at 37^*◦*^C with 1000 rpm shaking for exactly 30 min. The reaction was stopped with 30 *µ*l of EDTA, spun at 2000g for 5 min at 4^*◦*^C and resuspended in 100 *µ*l of PBS with 0.5% BSA and 0.1% Triton X-100.

##### FACS

Flow cytometry staining was carried out as described in “Flow cytometry of intracellular markers”. Samples processed in a BD Fusion cell sorter (Beckton Dickinson). 15,000 cells of each desired population were sorted in to 0.5ml of PBS with 0.5% BSA. Sorted cells were spun for 10 min at 3000 g and 4^*◦*^C and resuspended in Reverse crosslink buffer (50mM Tris-HCl ph8, 0.5% Tween-20, 0.5% Igepal CA-630) supplemented with 1 *µ*l of 20 mg/ml Proteinase K (Ambion, Cat. No. AM2546) and incubated overnight at 65^*◦*^C shaking at 300 rpm.

##### DNA isolation

The following day samples were spun down to collect condensation and DNA was isolated using the DNA clean up & Concentrator kit (Zymo, Cat. No. D4013) as per manufacturer’s instructions. DNA was eluted in 21 *µ*l of DNAse-free water. DNA was stored at -20^*◦*^C until ready to be processed.

##### Library generation

Transposed DNA was first amplified with 2X PCR Master Mix NEB (Cat. No. M0541S), and 1 *µ*l of constant forward primer and indexed reversed primer (Table S2) (both at 25 uM) in a total reaction volume of 50 ul. The program was as follows: 5 min at 72^*◦*^C, 30 s at 98^*◦*^C, followed by 7 cycles of 30 s at 98^*◦*^C, 30 s at 63 ^*◦*^C and 60 s at 72 ^*◦*^C, with a final extension of 5 min at 72 ^*◦*^C. DNA was cleaned up with 1.8x volumes of AMPureXP beads (Beckman Coulter, Cat. No. A63882) and eluted in 18 *µ*l of EB buffer. qPCR was carried out to determine the number of extra cycles using 2 *µ*l of the amplified DNA in technical duplicates. Each 20 *µ*l reaction contained 20 *µ*l of 2X SYBR Green PCR assay, using PowerUP SYBR Green Master Mix (Thermo Fisher, Cat. No. A25742) and 2.5 *µ*l of each primer (at 25 *µ*M each). The qPCR program was as follows: initial activation for 2 min at 50^*◦*^C, 30 s at 98^*◦*^C, followed by 40 cycles of 10 s at 98^*◦*^C, 30 s at 63^*◦*^C and 60 s at 72^*◦*^C. The number of additional cycles was calculated as 1/4 of the maximum amplification (Buenrostro et al., 2015). The second amplification was done using 12.5 *µ*l of amplified DNA, 2X PCR Master Mix NEB and 2.5 *µ*l of each primer (at 25 uM) in a total volume of 50 ul. The program was the same as the first amplification without the initial extension and for the calculated number of cycles. Finally, re-amplified DNA was cleaned up using 1.8x volumes AMPureXP beads and eluted in 30 ul. Samples were quantified in the QuBit, size profiles examined in the Bioanalyzer using the DNA High Sensitivity DNA Kit (Agilent, Part No. 5067-4626). Samples were pooled and sequenced in a NovaSeq by Novogene (Cambridge, UK).

#### Bulk ATAC-seq

##### Tn5 production

Methods were adapted from (Hennig et al., 2018) as follows: The pETM11-Sumo3-Tn5_E54K,L372P_ plasmid was obtained from EMBL-Heidelberg and transformed into *Escherichia coli* BL21 (DE3) Gold cells (Agilent). Bacterial cultures were grown at 30^*◦*^C to a density of OD_600_=0.6-0.8 and protein expression was induced with 0.5 mM isopropyl-*β*-D-thiogalactoside (IPTG). Prior to induction, cultures were cooled and maintained at 20^*◦*^C for protein expression overnight. Cell pellets were harvested by centrifugation for 10 minutes at 4000 rpm and then resuspended in lysis buffer (50 mM Tris-HCl pH 8.0, 20mM Imidazole, 0.5 M NaCl, 10% Glycerol, 1mM TCEP, 1U/ml Benzonase, 1 tablet/50ml Protease inhibitor tablets (cOmplete, Roche) and lysed with sonication. The cell lysate was then centrifuged for 30 minutes at 80,000 x *g*. The supernatant was collected and applied onto a 5ml Ni-Sepharose column (Cytiva). The column was washed with 10 Column Volumes wash buffer A (50mM Tris-HCl pH 8.0, 20 mM Imidazole, 0.5 M NaCl,10% Glycerol, 1mM TCEP). Bound proteins were eluted with buffer A containing 300mM Imidazole. Fractions containing His-Sumo3-Tn5_E54K,L372P_ were pooled and incubated overnight at 4^*◦*^C with SenP2 protease to remove the fusion tag. Next day, the sample was diluted six times and applied onto a 5ml Heparin HP column preequilibrated in buffer B (50mM Tris-HCl pH 8.0, 0.05 M NaCl, 1mM TCEP). Bound Tn5 protein was eluted with a linear gradient to 100% buffer B containing 1M NaCl, concentrated and loaded onto a Superdex 200 Increase 10/300 GL column (Cytiva) equilibrated in buffer C (50 mM Tris pH 8.0, 150 mM NaCl, 1 mM TCEP and 5% glycerol). Fractions corresponding to the Tn5 peak were pooled, concentrated to 10*µ*M and flash frozen in liquid nitrogen.

##### Tn5 assembly

Dilution and oligo assembly of Tn5 was performed as previously described (Corces et al., 2017; Ma et al., 2020). In brief: purified protein was diluted to 4 *µ*M in Dilution Buffer (50 mM Tris, 100 mM NaCl, 0.1 mM EDTA, 1 mM DTT, 0.1% NP-40, and 50% glycerol). Two independent oligo mixes were assembled (Mix A, Mix B), each with 5 *µ*l of 100 *µ*M Tn5MEREV oligo, 5 *µ*l of 100 *µ*M of either Tn5 1 (Mix A) or Tn5 2 ME comp (Mix B), and 40*µ*l of nuclease-free water by incubating each mix in a PCR thermocycler as follows: 95^*◦*^C for 3 minutes, 65^*◦*^C for 3 minutes and ramp down to 24^*◦*^C at a rate of -1C/min. Once annealed, Mix A and Mix B were combined with 100 *µ*l glycerol to create a 5 *µ*M, 50% glycerol adaptor mix. Equal parts of diluted Tn5 transposase and adaptor mix were mixed and incubated at 25^*◦*^C for 60 min. For oligo sequence see (Table S2)

##### Transposition and library generation

Wells were washed and cells were dissociated using 0.5 ml Accutase per well of a 6 well plate incubated 5 min at 37^*◦*^C. Cells were washed in PBS, and 50,000 cells per condition and replicates were used for ATAC-seq following established protocols (Corces et al., 2017; Yoshida et al., 2019) (for sequences used see (Table S2)) and as described for CaTS-ATAC. After transposition, DNA was directly purified and used for library preparation (as described for CaTS-ATAC).

#### Cell type specific RNA-seq based on intracellular markers

Samples for CaTS-RNAseq were collected in the same way as for CaTS-ATAC but all buffers were supplemented with 1 *µ*L/ml of RNasin Plus (Promega, Cat. No. N2615). After fixation and quenching, samples were stained for intracellular TFs, also as described in “Flow cytometry of intracellular markers” with RNasin-supplemented buffers.

Cells were sorted into 0.5 ml PBS with 0.5% BSA and 1 *µ*l/ml of RNasin Plus. RNA extraction from fixed samples were performed with the RecoverAll RNA/DNA Isolation Kit (Thermo Fisher Scientific, Cat. No. AM1975) with the modifications described in (Amamoto elife). In brief, sorted cells were spun down at 3000 g for 7 min at 4^*◦*^C and as much supernatant as possible was removed (leaving 40-50 *µ*l). 100 *µ*l of Digestion Buffer previously mixed with 4 *µ*l of Protease mix was added to each sample and resuspended. The tubes were incubated at 50^*◦*^C for 3 h and then stored at -80^*◦*^C. RNA isolation was performed as per manufacturer’s instructions, eluting in 17 *µ*l of Nuclease-free water.

Due to low quality and limiting amounts, the maximuM amount (10.5 *µ*l) was used for library preparation using the SMART-Seq HT kit (Takara, Cat. No. 634437) followed by Nextera XT DNA Library Preparation Kit (Illumina, Cat. No. FC-131-1096).

#### ATAC-seq processing

Data was processed using the nf-core atacseq pipeline (https://nf-co.re/atacseq) with the following options: --genome mm10 --macs fdr 0.00001 --skip diff analysis --min reps consensus 2 -r 1.2.0. Peaks were further filtered out if they did not pass qval < 0.00001 in at least 1 sample (qval generated from MACS peak calling in each sample).

#### ATAC-seq differential expression analysis

Read counts within the consensus intervals generated by featureCounts were used as input for DESeq2 (ref Love). Principal Component Analysis was performed using the top 30000 most variable elements and colored by different sample metatada.

To assess the cell type specific accessibility for the same NP cell types at different SAG concentrations, all by all differential expression was performed for all conditions at day 5. Number of differentially accessible intervals plotted (fold change > 2, basemean > 100) as a bar graph, and individual elements are plotted color-coded by their p-adjusted values for specific pairwise comparisons.

Selection of differentially accessible elements across cell types and timepoint was performed by pairwise DESeq2 analysis between any two cell type within a timepoint and any two timepoints for the same cell type. Elements that fulfilled padj < 0.01 & abs(log2FoldChange) > 2 & baseMean > 100 were selected for subsequent clustering.

#### ATAC element clusters by kmeans

Varianced stabilized transformed data generated using DESeq2 were used as input to identify clusters of elements with the same dynamics. Clustering was performed using kmeans with a high number of centers, 30, and subsequently re-grouping clusters of very similar dynamics using hclust and target of 9 final clusters. This was chosen as indepent iterations resulted in reproducible clusters and dynamics.

#### Analysis of published ChIP-seq data

Processing of published ChIP-seq datasets was for NKX6.1, NKX2.2, OLIG2 (Nishi et al., 2015), SOX2 and FOXA2 (Peterson et al., 2012) from neural embryoids with SAG were processing using the nf-core chipseq pipeline as follows: nextflow run nf-core/chipseq --input design.csv --single end --genome mm10 -profile crick -r 1.1.0.

FOXA2 from endoderm differentiations were process with the nf-core atacseq pipeline due to lack of input samples. nextflow run nf-core/atacseq --input design.csv --single end --mito name false --genome mm10 -profile crick -r 1.1.0.

#### Footprinting analysis

BAM files for merged replicates were used as input for TOBIAS ATACorrect (Bentsen et al., 2020), followed by TOBIAS Footprint. TOBIAS BINDetect was run on all samples combined using the following the JASPAR2018, HOCOMOCO and Taipale databases (Jolma et al., 2013; Khan et al., 2017; Kulakovskiy et al., 2017).

Motifs that were amongst the top 5% highest absolute fold change or 5% smallest pvalues in each pair-wise comparison were selected. Motifs were grouped in archetypes (Vierstra et al., 2020). The most variable archetypes for the desired comparison (either p0-1, p2, pM at day 5 and 6, or all conditions) were selected and RNA expression for each TF that is associated with the archetype was compared to the motif score for that archetype. This correlation analysis between motif score and RNA expression was used to find candidate TFs driving the chromatin changes observed.

#### RNA-seq data processing

Data was processed using the nf-core rnaseq pipeline with the following options: --star index ‘star ercc mm10 genome’ --gtf ―mm10.refGene wERCC.gtf’ --fc group features type gene id --fc extra attributes gene id -r 1.4.2. Gene counts were obtained from featureCounts with the option ignoreDup=TRUE to try and remove PCR duplicates. Samples were excluded from downstream analysis based on a number of QC criteria particularly applicable to this type of low input samples from PFA-fixed cells including: too small proportion of the library being of mouse origin (excess spike-in representation), large number of overepresented sequences, and loss of dynamic range in spike-in quantification.

## Data and software availability

The accession numbers for the data generated in this paper (CaTS-ATAC, RNA-seq and bulk ATAC-seq) is GSE204921.

## Code availability

All analysis scripts are available at https://github.com/MJDelas/Neural_DV_ATAC

## I. SUPPLEMENTARY MATERIAL

**FIG. S1:**
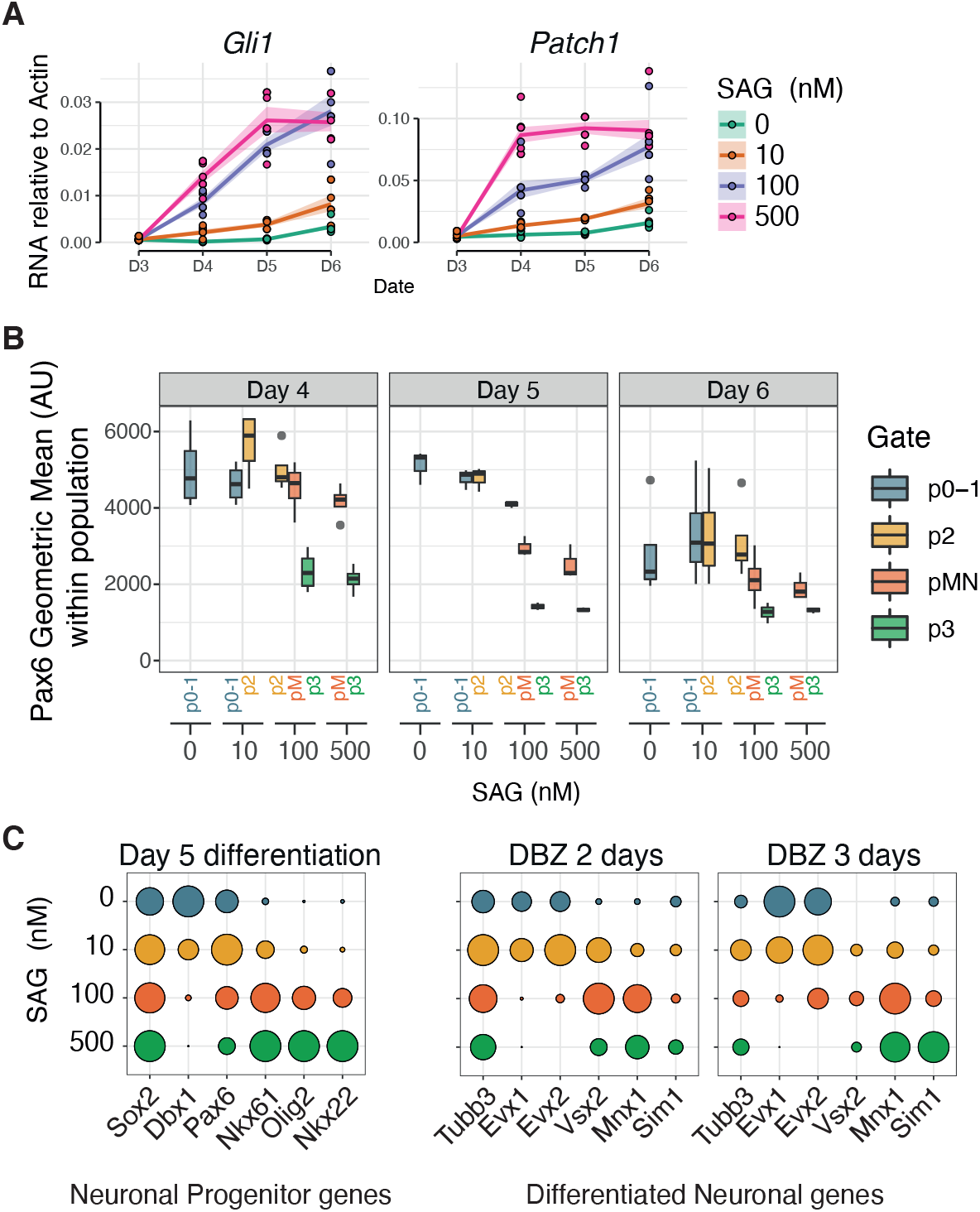
A cellular model of ventral spinal cord neural progenitors recapitulates aspects of in vivo patterning, Related to Figure 1. (A) Relative expression (RT-qPCR) for Shh responsive genes show a SAG concentration-dependent response. (B) Pax6 levels recapitulate the lower expression in pMN compared to p0-1 and p2 observed for this protein in vivo. Pax6 is not expressed in p3 NPs. (C) Relative expression (RT-qPCR) for progenitors and neuronal markers at Day 5 (predominantly progenitors) and 2 or 3 days after neuronal induction with Notch inhibitor dibenzazepine (DBZ) show the expected enrichment for dorsoventral progenitors and neuronal markers as a function of SAG concentration.

**FIG. S2:**
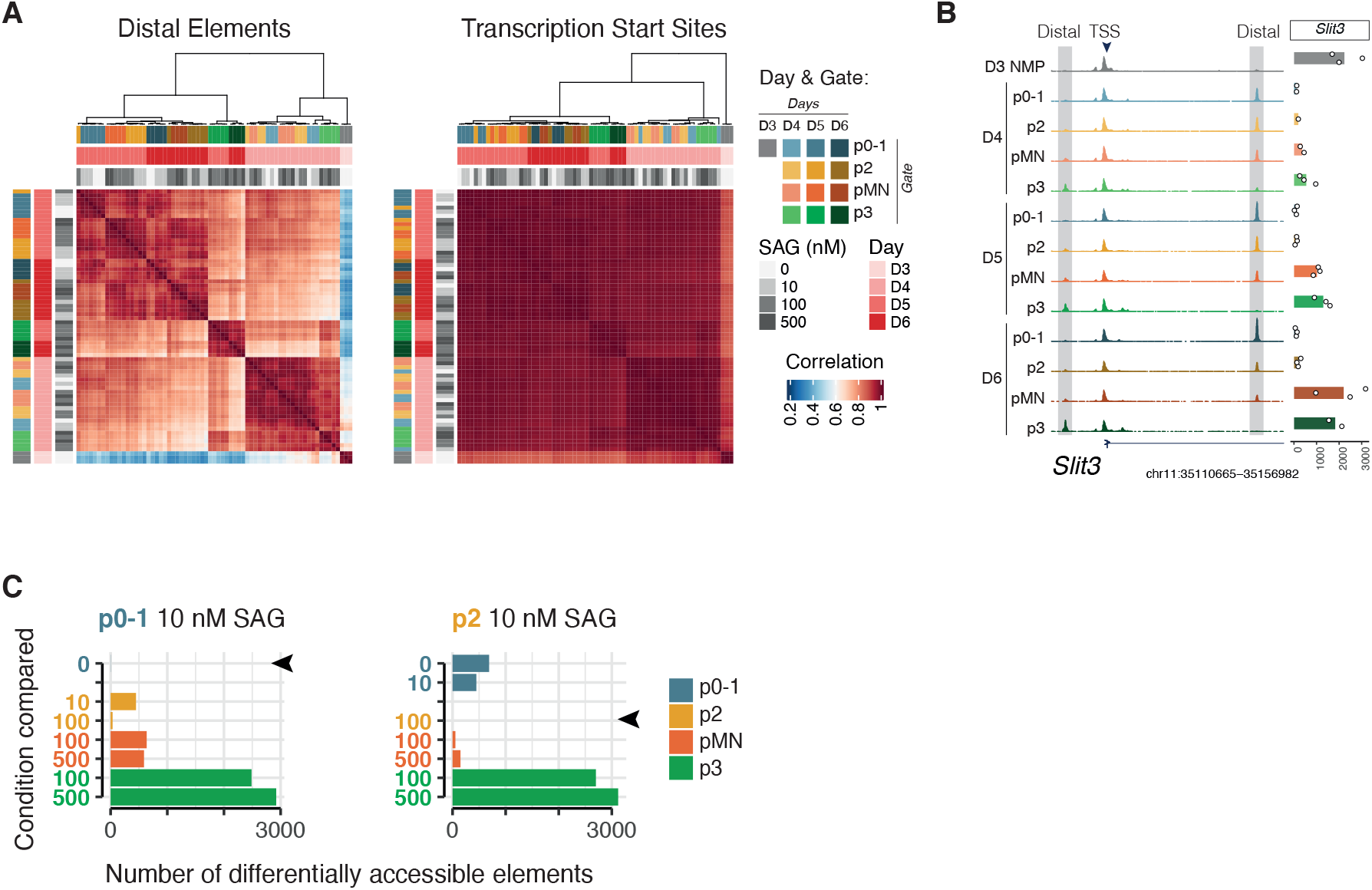
Cell type specific ATAC-seq characterization, Related to Figure 2. (A) Sample to sample correlation based on distal elements shows sample clustering by cell type. Correlation based on TSS is less cell type specific. (B) Example locus showing broad accessibility across the promoter in all samples and cell type specific variations in accessibility at distal elements. (C) Number of differentially accessible elements between the indicated samples and all other conditions reveals few or no differentially accessible elements between the same cell type from a different concentration of SAG.

**FIG. S3:**
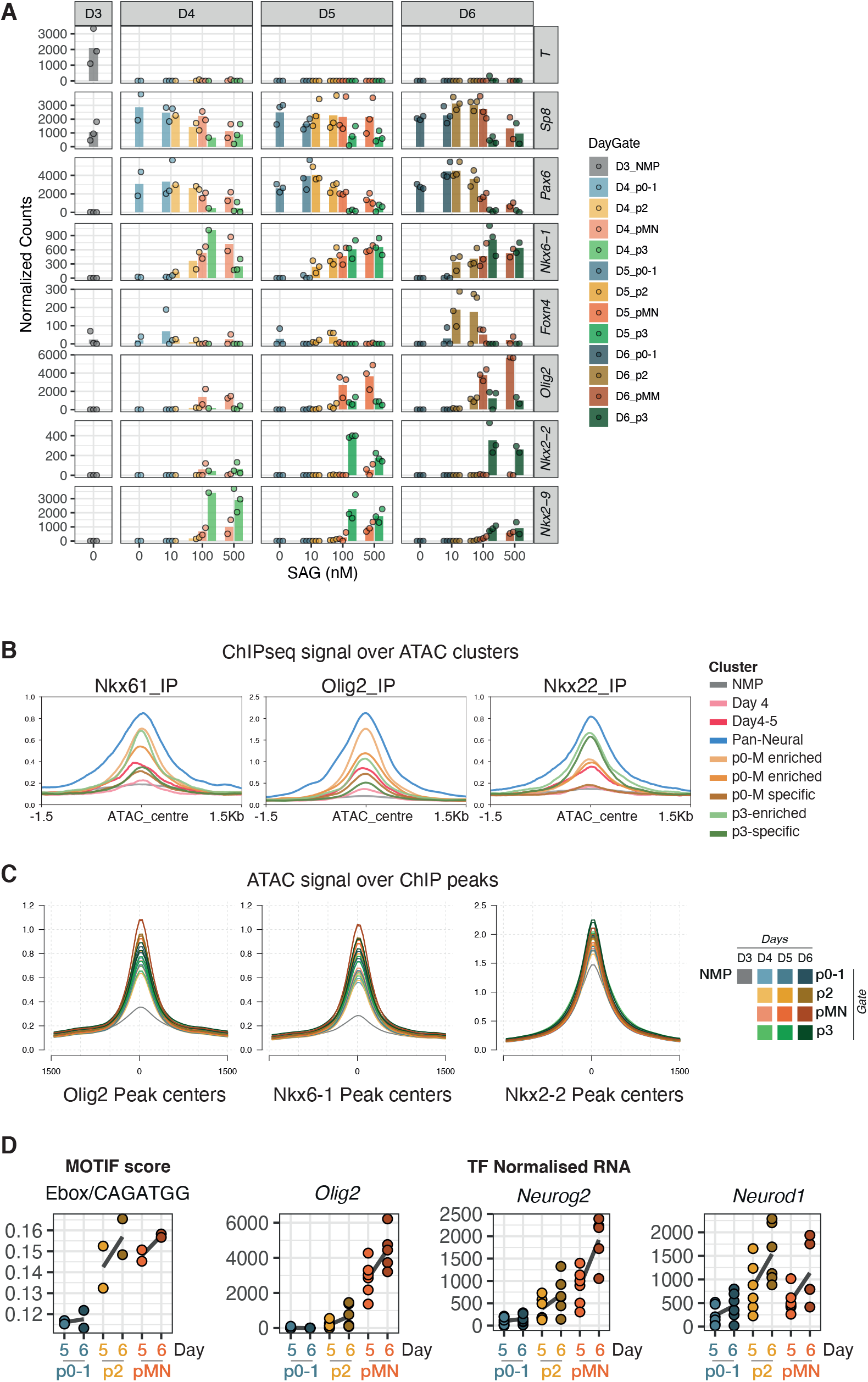
A shared accessibility landscape for p01-, p2 and pM, Related to Figure 3. (A) RNA-seq normalized gene counts for key marker genes validates cell type identity of sorted populations. (B) Normalized ChIP-seq coverage of each ChIP-seq data set over the groups of elements belonging to each ATAC clusters from Fig 3B shows binding of all TFs to pan-neural and cell type specific clusters. (C) Normalized accessibility coverage of each sample over the peak called regions for each ChIPseq data set shows accessibility of all NPs over the regions bound by all TFs. (D) Footprinting score for the archetype Ebox/CAGATGG is higher in p2 and pM. Normalized RNAseq expression for the top correlated three genes associated with this motif reveals their potential contribution to the footprint signal in different cell types.

**FIG. S4:**
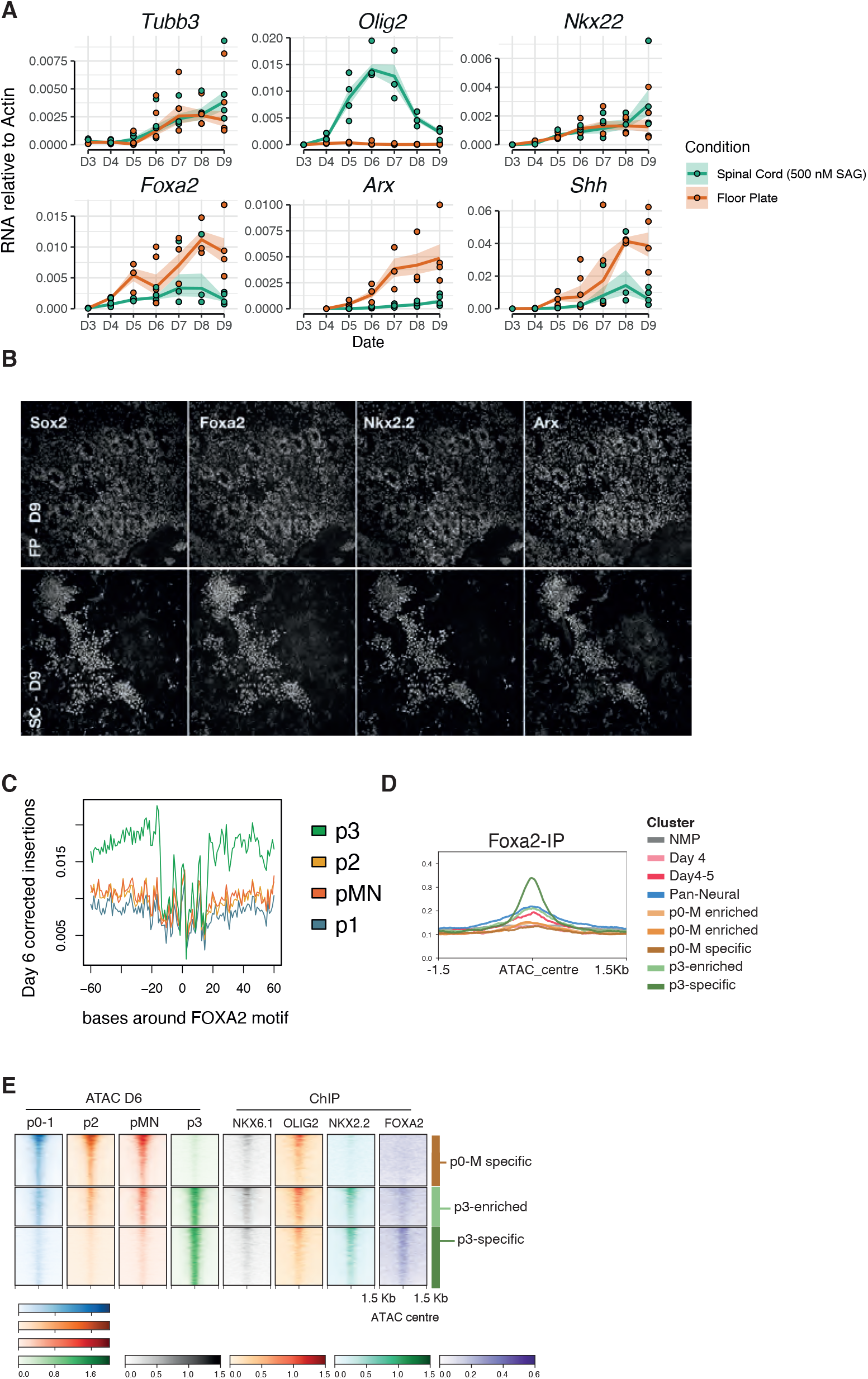
Spinal cord neural differentiation generates minimal amounts of floor plate and Foxa2 binding p3 NPs, Related to Figure 4. (A) Relative mRNA (RT-qPCR) for the genes indicates shows clear induction of floor plate (FP) markers in FP conditions and minimal induction in spinal cord neural differentiations. (B) Immunohistochemistry staining for shows robust induction of FP markers in FP conditions. (C) Representative metaplot of corrected insertions over the FOXA2 motif across all accessible regions for all NPs at Day 6 shows a footprint in p3 NPs. (D) Normalized FOXA2 ChIP-seq coverage over the groups of ATAC-seq elements in each cluster from Fig 3B shows accessibility in p3-specific elements. (E) Coverage heatmap for ATAC and ChIP-seq for the p0-M specific, the p3-enriched and the p3-specific cluster shows binding of NKX2.2 and FOXA2 to p3-specific sites.

**FIG. S5:**
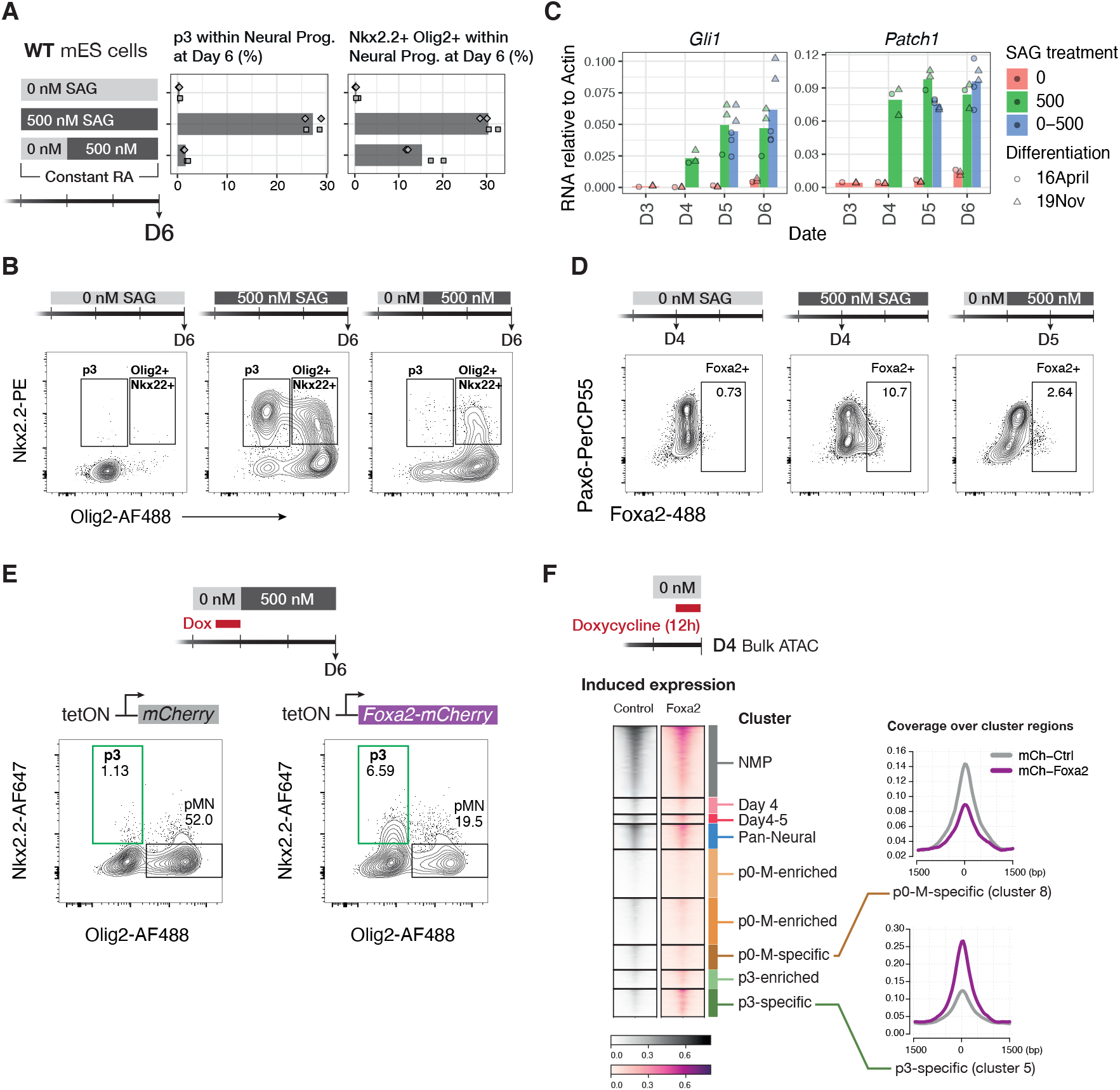
Delayed SAG abolishes p3 generation due to lack of Foxa2 induction, Related to Figure 5. (A) Generation of p3 at day 6 is greatly impaired if SAG administration is delayed by 24h (left graph). Differentiations do express Nkx2.2 but co-expressed with Olig2 (right). (B) Representative flow cytometry plots for the data quantified in (A). Gated on SOX2+ live neural progenitors. (C) Relative expression of Shh target genes shows cells respond to signalling in the delayed SAG regime (“0-500”). (D) Flow cytometry of Foxa2 intracellular antibody staining without SAG administration of 24h after SAG administration shows reduced Foxa2 induction in the SAG delayed regime. (E) Doxycycline-induced expression of Foxa2 for 12h in the SAG delayed regime rescues p3 generation at day 6. (F) Bulk ATAC seq coverage after overexpression of Foxa2 or control for the ATAC clusters identified in Fig 3B. (G) Metaplot of normalized coverage comparing ATAC in Foxa2 and control overexpressed for cluster p0-M-specific and p3-specific show increased accessibility of Foxa2 overexpression in p3-specific sites.

**FIG. S6:**
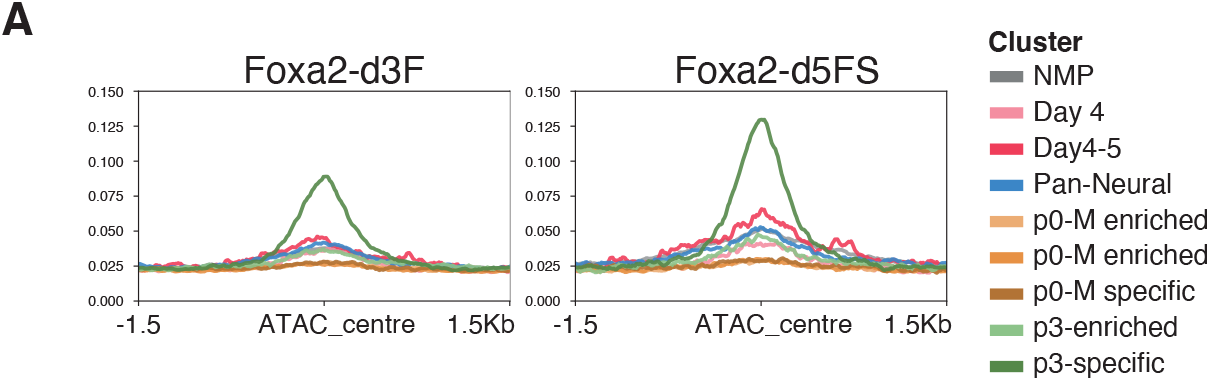
Binding of endodermal Foxa2 to ventral neural sites, Related to Figure 6. (A) Normalized Foxa2 ChIP-seq coverage over the ATAC-seq clusters identified in Fig 3B shows accessibility in p3-specific sites.

**TABLE S1:** Antibodies used in this study

| **Antibody** | **Source** | **Cat. No.** |
| --- | --- | --- |
| Sox2-V450 | BD Biosciences | 561610; clone O30-678 |
| Pax6-PerCPCy5.5 | BD Biosciences | 562388; clone 018-1330 |
| Nkx6.1-AlexaFluor647 | BD Biosciences | 563338; clone R11-560 |
| Goat anti-Olig2 | R&D | AF2418 |
| Nkx2.2-PE | BD Biosciences | 564730; clone 74.5A5 |
| Tubb3-Biotin | BD Biosciences | 560394; TUJ1 |
| Foxa2- AlexaFluor488 | SC Biotechnology | sc-374376 AF488; clone H-4 |
| Goat anti-Sox2 | R&D | AF2018 |
| Mouse anti-Nkx2.2 | BD Biosciences | 564731, clone 74.5A5 |
| Rabbit anti-Olig2 | Merck Millipore | AB9610 |
| Rabbit anti-Pax6 | Merck Millipore | AB2237 |
| Donkey anti-mouse AF488 | Invitrogen | A21202 |
| Donkey anti-goat AF488 | Invitrogen | A11055 |
| Donkey anti-rabbit AF488 | Invitrogen | A21206 |
| Strep-APC-Cy7 | BioLegend | 405208 |
| Donkey anti-mouse AF568 | Invitrogen | A10037 |
| Donkey anti-goat AF568 | Invitrogen | A11057 |
| Donkey anti-mouse AF647 | Invitrogen | A31571 |
| Donkey anti-goat AF647 | Invitrogen | A21447 |

TABLE S2: Oligonucleotides used in this study (see supplementary material)

## Notes

https://www.ncbi.nlm.nih.gov/geo/query/acc.cgi?acc=GSE204921

https://github.com/MJDelas/Neural_DV_ATAC

### Competing Interest Statement

The authors have declared no competing interest.

